# Limitation by a shared mutualist promotes coexistence of multiple competing partners

**DOI:** 10.1101/2020.04.22.055517

**Authors:** Sarah P. Hammarlund, Tomáš Gedeon, Ross P. Carlson, William Harcombe

**Author notes:** Email: (SPH), (WH).

## Abstract

Although mutualisms are often studied as simple pairwise interactions, they typically involve complex networks of interacting species. How multiple mutualistic partners that provide the same service and compete for resources are maintained in mutualistic networks is an open question. We use a model bacterial community in which multiple ‘partner strains’ of *Escherichia coli* compete for a carbon source and exchange resources with a ‘shared mutualist’ strain of *Salmonella enterica*. In laboratory experiments, competing *E. coli* strains readily coexist in the presence of *S. enterica*, despite differences in their competitive abilities. We use ecological modeling to demonstrate that a shared mutualist can create temporary resource niche differentiation by limiting growth rates, even if yield is set by a resource external to a mutualism. This mechanism can extend to maintain multiple competing partner species. Our results improve our understanding of complex mutualistic communities and aid efforts to design stable microbial communities.

## Introduction

Mutualisms—bidirectional positive interspecies interactions—are abundant and important^1,2^. Traditionally, studies of mutualism have focused on interactions between two species. However, communities often contain many species of mutualists that interact in complex networks^3,4^. For example, many flowering plants are pollinated by multiple insect species^5^, and corals interact with a phylogenetically diverse set of endosymbionts^6^. In fact, two-partner mutualisms are now thought to be the exception rather than the norm^3,7^. Understanding the ecology of multiple mutualist communities is an important goal^3,8^. To do so, we need theoretical predictions and experimentally tractable multiple mutualist communities.

In many multiple mutualist systems, several functionally-similar species within a “partner guild” supply resources or services to a “shared mutualist,” which supplies resources in return (**Fig. 1**)^3,8^. For example, a guild consisting of pollinators like bees and butterflies may exchange pollination services with a shared plant host that provides both pollen, from which bees benefit, and nectar, from which butterflies benefit^8^. Recent studies have suggested that interactions between species within the partner guild can affect coexistence and stability of the whole community^9,10^. Within-guild interactions may be especially important if the partner mutualists within the guild are ecologically similar. If partner species’ resource niches overlap, they may compete for resources that are external to the mutualism. When multiple species compete for the same limiting resource, one species may competitively exclude the others, leading to a loss of diversity within the community^10,11^. However, because species-rich communities of multiple mutualists exist in nature, certain mechanisms that maintain coexistence must exist. Here, we seek to understand the conditions in which multiple partner mutualists are able to coexist despite competition for a common resource.

**Figure 1.**
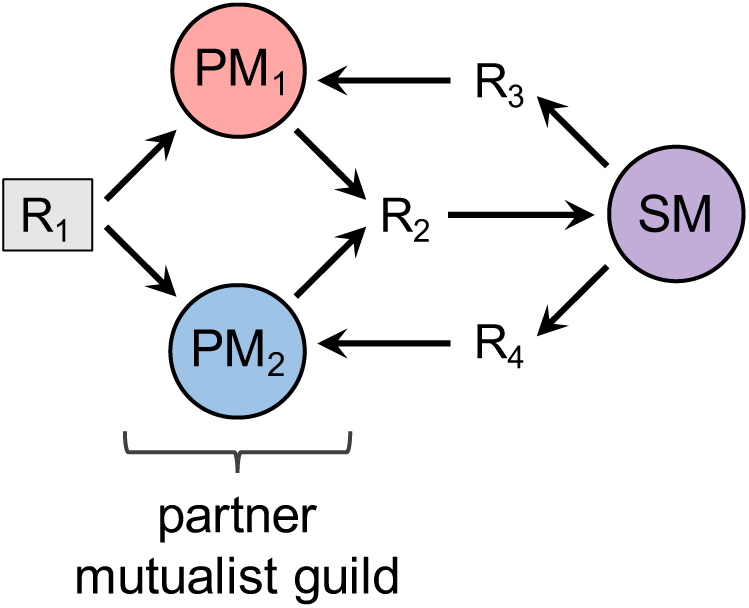
A multiple mutualist system with resource competition between partner species. The partner mutualist guild is composed of two partner mutualists (PM_1_ and PM_2_) that compete for access to resource R_1_ (shaded), and produce resource R_2_. The shared mutualist (SM) consumes R_2_ and produces two different resources, R_3_ and R_4_. R_3_ is consumed by PM_1_ and R_4_ is consumed by PM_2_. An example of such a community is a shared mutualist plant that provides pollen to a bee (PM_1_) and nectar to a butterfly (PM_2_). The bees and butterflies may compete for other nutrients that are external to the mutualism.

Using a community of mutualistic bacteria, we explore the potential for coexistence of multiple partner species. Our system consists of a partner guild of *Escherichia coli* strains that compete with one another for a carbon source, and engage in mutualism with a strain of *Salmonella enterica*, the shared mutualist (**Fig. 2a**). The strains engage in mutualism via cross-feeding, with the *E. coli* strains providing acetate and receiving amino acids from *S. enterica*. We show that the competing *E. coli* strains are unable to coexist when they are provided amino acids in the growth media rather than obtaining them from the shared mutualist, because one strain has a faster growth rate. However, when the shared mutualist is added to the community, the two *E. coli* strains coexist, maintaining the diversity of the multiple mutualist community. Next, we use a resource-explicit ecological model to identify factors that promote coexistence. We show that limitation by the shared mutualist is key— if the shared mutualist sets the growth rate of the community, the two partner mutualists coexist because they are temporarily limited by different resources. Finally, we demonstrate computationally that this phenomenon can promote the coexistence of more than two partner mutualists. This work helps us understand how diversity is maintained in multiple mutualist communities and can inform efforts to design stable microbial communities.

**Figure 2.**
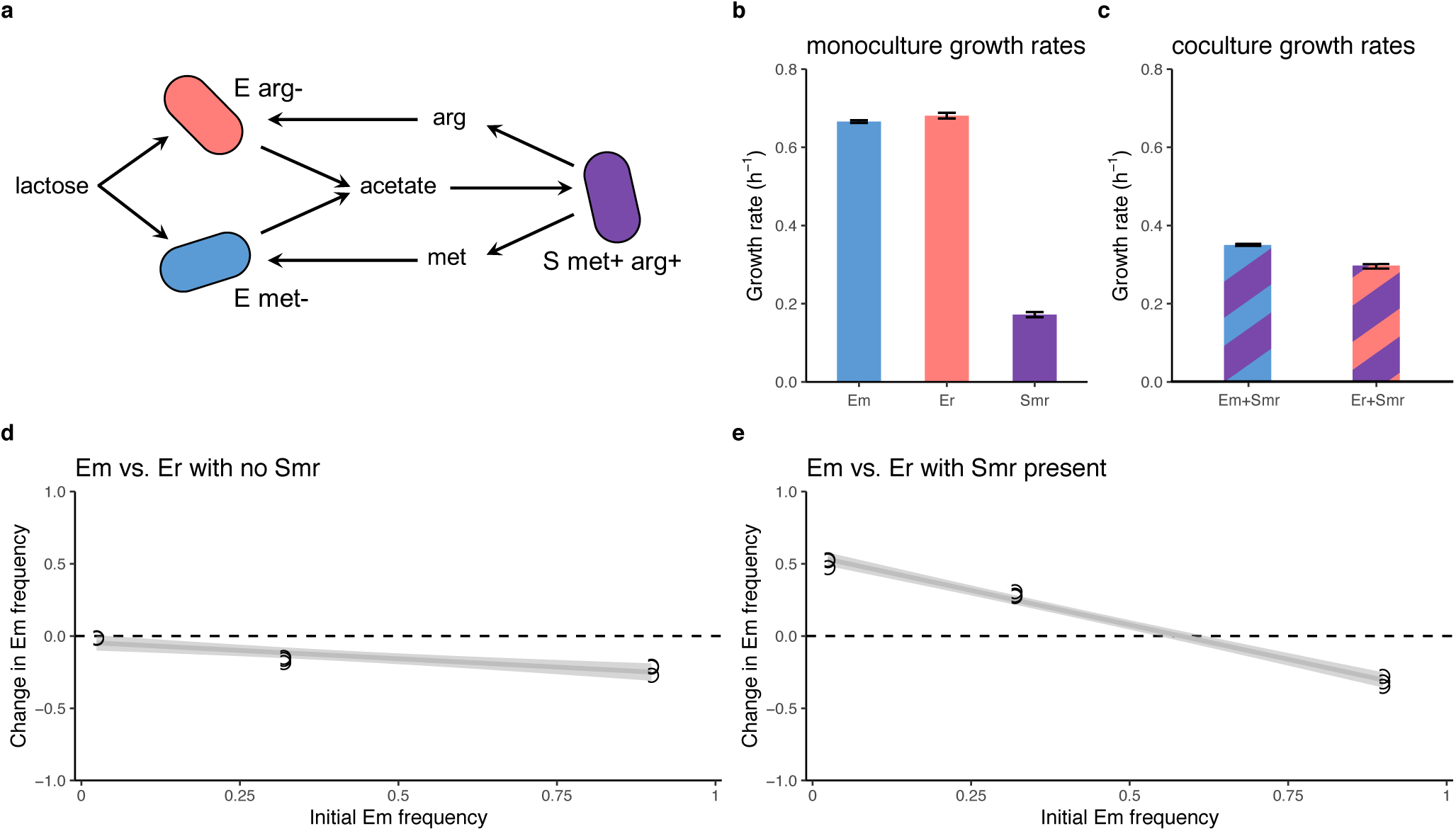
Coexistence in a multiple mutualist community of cross-feeding bacteria. **a**, Schematic showing the interactions between the *E. coli* partner mutualists that comprise the partner guild and the *S. enterica* shared mutualist. The arginine auxotroph, E arg- (“Er”), and the methionine auxotroph, E met- (“Em”), consume lactose and produce acetate. S met+ arg+ (“Smr”) consumes acetate and produces both arginine (arg) and methionine (met). **b**, The growth rates in monoculture of the three strains differ (one-way ANOVA: F(2, 9) = 9897, p < 1e-15). Er grows faster than Em (Tukey HSD: p = 0.037), and Smr grows much more slowly than either *E. coli* strain (Tukey HSD: p < 1e-7). Em and Er were grown in lactose media with excess methionine and arginine, and Smr was grown in acetate media. Error bars represent +/- one standard deviation. **c**, Growth rates of Em+Smr and Er+Smr cocultures. Em+Smr cultures grow faster (two-tailed t-test: df = 2, p-value = 0.004). Error bars represent +/- one standard deviation. **d**, A mutual invasibility experiment with cocultures of Em and Er in a lactose medium with excess amino acids. The frequency of Em decreases for all starting frequencies, including when started at 2% of the population (two-tailed t-test: df = 2, p = 0.005), indicating that Em is the weaker competitor for lactose. The change in Em frequency was calculated as [Em/ (Em + Er)]_final_ – [Em/(Em + Er)]_initial_. **e**, A mutual invasibility experiment in the three-strain multiple mutualist community in a lactose medium. Em increases in frequency when started rare (two-tailed t-test: df = 2, p = 0.001), but decreases in frequency when started common (two-tailed t-test: df = 2, p = 0.004), indicating that the two *E. coli* strains coexist. The change in Em frequency was calculated as [Em/ (Em + Er)]_final_ – [Em/(Em + Er)]_initial_. Smr yields are similar across all three Em frequencies (**Fig. S4**).

## Results

### Laboratory experiments

We studied competition between two partner mutualists using a laboratory system of cross-feeding bacteria (**Fig. 2a**). The partner guild consists of one *E. coli* strain that is a methionine auxotroph (“Em”) and another *E. coli* strain that is an arginine auxotroph (“Er”)—each strain lacks a gene in the biosynthetic pathway for its respective amino acid, so in order for a strain to grow, its required amino acid must be available in the environment. The two *E. coli* strains compete for lactose, which we provide in the growth media, and excrete acetate as a byproduct of lactose metabolism. We experimentally evolved a “shared mutualist” strain of *Salmonella enterica* (“Smr”) that secretes methionine and arginine. Smr was derived from a strain that we had previously evolved to secrete methionine^12^, and acquired a mutation in *argG* causing arginine secretion. Smr consumes acetate, and is unable to metabolize lactose.

Classically, species cannot coexist if they have different growth rates and compete for the same limiting resource^11^. Therefore, we started by measuring the growth rates of the three strains in monoculture and in pairwise coculture. For all experiments, we used a batch culture setup, in which populations grow until resources are depleted. When grown in monoculture in media containing each strain’s required nutrients, the three strains have different maximum growth rates (one-way ANOVA: F(2, 9) = 9897, p < 1e-15; **Fig. 2b**). Er has a slightly higher growth rate than Em (Tukey HSD: p = 0.037), and both *E. coli* strains grow faster than Smr (Tukey HSD: p < 1e-7 for both comparisons). When each *E. coli* strain is grown separately in coculture with Smr in lactose media with no amino acids, the coculture containing Em grows faster (two-tailed t-test: df = 2, p = 0.004; **Fig. 2c**). This may be because Smr secretes methionine at a faster rate than arginine, or because Em requires less methionine than Er requires arginine (**Fig. S1**). Yields in monoculture and coculture are shown in **Fig. S2** and **Fig. S3**.

Next, we tested whether the two *E. coli* strains coexist in a lactose environment with excess amino acids and no Smr present. We predicted that the strain with the faster monoculture growth rate, Er, would outcompete Em. We assessed coexistence of the two *E. coli* strains through a mutual invasibility test, measuring whether each *E. coli* strain could increase in frequency when initially rare. Mutual invasibility would indicate negative frequency dependence and coexistence^13,14^. Em decreased in frequency from all three initial frequencies, indicating that Er outcompetes Em and coexistence is not possible (**Fig. 2d**; two-tailed t-test for the lowest initial frequency: df = 2, p = 0.005).

These findings led to two alternative hypotheses about coexistence in the three-strain community: Hypothesis 1) One *E. coli* strain will outcompete the other. Er may outcompete Em due to its faster monoculture growth rate and greater competitive ability, or Em may outcompete Er due to its faster coculture growth rate when paired with Smr. Hypothesis 2) The two *E. coli* strains coexist.

To assess coexistence of the two *E. coli* strains in the three-strain community, we again conducted a mutual invasibility test. We inoculated cultures with three different initial frequencies of the *E. coli* strains, with a constant initial population size of Smr. In line with Hypothesis 2, both *E. coli* strains increased in frequency when initially rare, indicating coexistence (**Fig. 2e**). Em increased in frequency when initially rare (two-tailed t-test: df = 2, p = 0.004), and decreased in frequency when initially common (two-tailed t-test: df = 2, p = 0.001). We have previously shown that *S. enterica* coexists with a single strain of cross-feeding *E. coli* through similar mutual invasibility experiments^15^. Yields are shown in **Fig. S4**.

### Ecological modeling

To understand why the two *E. coli* strains coexist, we constructed an ecological model with ordinary differential equations for the three strains (Em, Er, and Smr) and the four resources (lactose, acetate, methionine, and arginine):

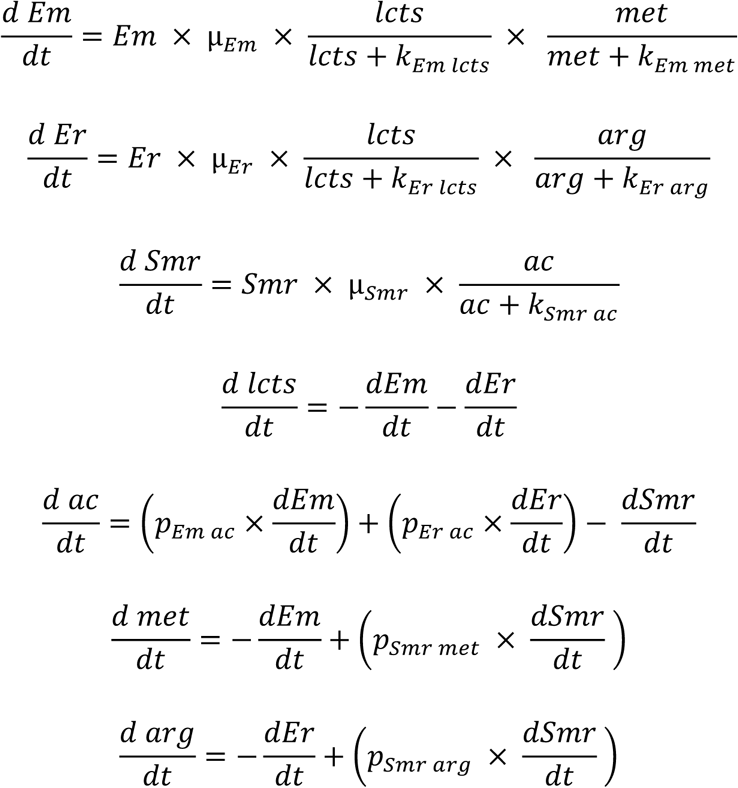

Em, Er, and Smr are the population densities of each strain (cells/ml). Resources (lcts = lactose, ac = acetate, met = methionine, arg = arginine) are in units of cell-equivalents/ml (the density of cells that a unit of resource can produce). Growth is governed by Monod saturation rates using Monod constants (e.g. k_Em lcts_), which are in units of cells/ml. Production terms (e.g. p_Em ac_) are in units of cells/cell. Default values and parameter descriptions can be found in **Table S1**. Briefly, we kept the model simple by using equal values for the same parameters for each of the three strains, except for their growth rates, which we approximated based on relative growth rates in the lab system. Default growth rates are µ_Em_ = 1.0, µ_Er_ = 1.1, and µ_Smr_ = 0.5 with units of 1/timestep.

Consistent with our lab system, Er outcompetes Em when the two are grown in an environment without Smr and with unlimited amino acids (**Fig. 3a**). This is because Er has a faster growth rate than Em. Also consistent with our findings in the lab system, in the three strain community with no amino acids provided, the two *E. coli* strains coexist. Both *E. coli* strains increase in frequency when started rare (**Fig. 3b**).

**Figure 3.**
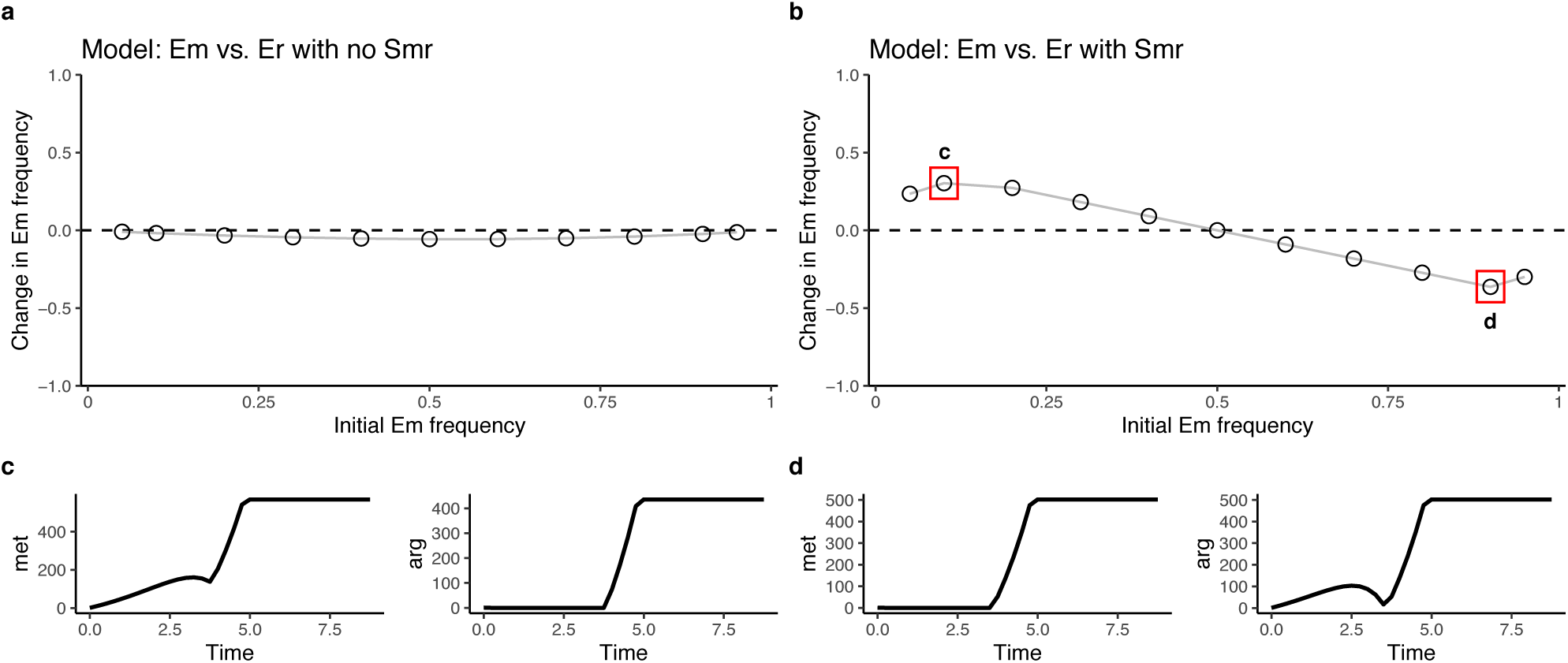
A resource-explicit model shows that temporary limitation by different resources promotes coexistence. **a**, In an Em+Er coculture in an environment with lactose, methionine, and arginine, Em decreases across a range of initial frequencies, indicating that Er is the stronger competitor and that the two strains cannot coexist. **b**, In a community of Em, Er and Smr in a lactose environment, both Em and Er are able to increase in frequency when initially rare, which indicates coexistence through negative frequency dependence. **c**, Dynamics of methionine and arginine at the left data point boxed in red in part b, where Em begins at 10%. During early timepoints, methionine is not limiting, while arginine is limiting. Growth ceases when lactose is depleted, and dynamics of the three strains and all resources are shown in **Fig. S5. d**, Dynamics of methionine and arginine at the right data point boxed in red in part b, where Em begins at 90%. During early timepoints, arginine is not limiting, while methionine is limiting. Dynamics of strains and all resources are shown in **Fig. S6**.

Examining the dynamics of amino acid concentrations provides a potential explanation for coexistence in the three-strain system. When Em starts rare (the left boxed point in **Fig. 3b**), there is plentiful methionine at all timepoints, while arginine is limiting during the start of growth (**Fig. 3c, Fig. S5**). Conversely, when Er starts rare (the right boxed point in **Fig. 3b**), arginine is never limiting, while methionine is limiting at early timepoints (**Fig. 3d, Fig. S6**). This means that the initially-common *E. coli* strain’s growth rate is limited by its amino acid, while the initially-rare *E. coli* strain is able to grow at its maximum growth rate, because its amino acid is abundant. The initially-rare *E. coli* strain is therefore able to increase in frequency.

To explore the importance of amino acid limitation for coexistence, we investigated the influence of Smr’s growth rate and amino acid production rates. We hypothesized that these parameters are key for coexistence because they affect amino acid limitation. In our lab system, Smr grows more slowly than both *E. coli* strains (**Fig. 2b**), but using our model, we can explore the effect of increasing Smr’s growth rate. We increased Smr’s growth rate from 0.5 to 1.5. To test for coexistence, we started Em rare (10%) and tracked its change in frequency. When Smr’s growth rate is lower than 0.96, Em increases in frequency and the two *E. coli* strains coexist (**Fig. 4a**). However, when Smr’s growth rate is greater than 0.96, Em decreases in frequency and Er takes over. Under these conditions, methionine and arginine are never limiting (compare **Fig. 4a** inset plots). This means that both *E. coli* strains grow at their maximum growth rates until lactose is depleted, and the strain with the faster growth rate takes over. Smr’s amino acid production rates also affect coexistence. We measured whether Em is able to increase in frequency from rare across a range of arginine production rates, keeping the methionine production rate fixed at 1 and Smr’s growth rate at 0.5. When Smr grows more slowly but produces arginine at a rate four times faster than methionine, Em is not able to invade from rare and there is no coexistence (**Fig. 4b**). This is because Er is no longer arginine-limited (amino acid dynamics are similar to inset plots in **Fig. 4a**).

**Figure 4.**
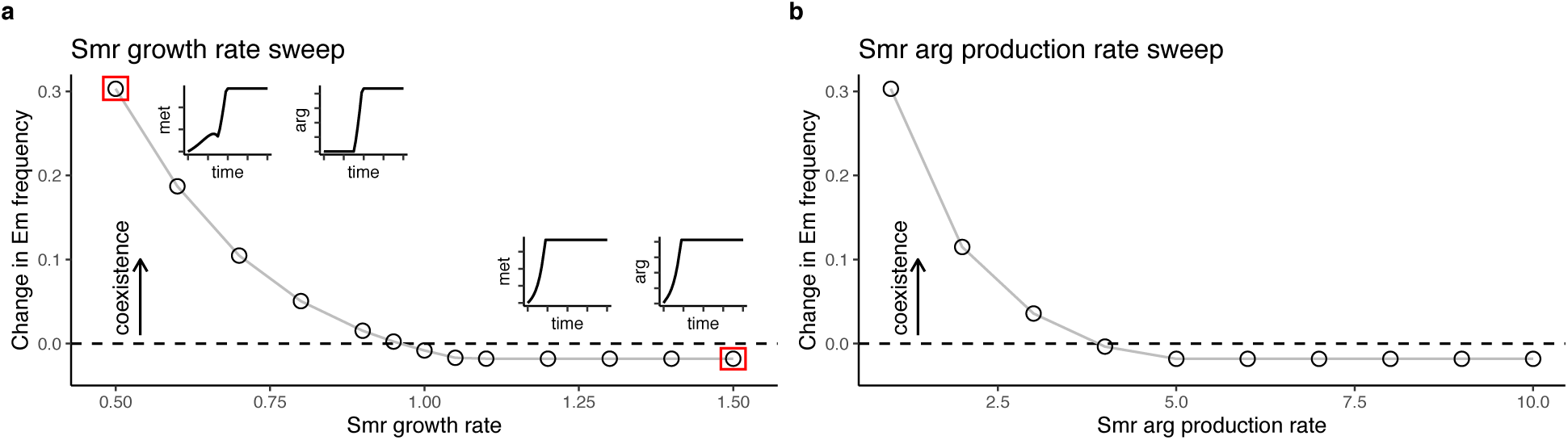
The shared mutualist’s growth rate and amino acid production rates affect coexistence. **a**, Smr growth rate affects coexistence. Growth rates of Em and Er are set at their default levels (µ_Em_ = 1 and µ_Er_ = 1.1) and coexistence is evaluated across a range of Smr growth rates. In these simulations, Em begins at a frequency of 0.1, and coexistence is indicated by an increase in frequency. Coexistence is possible when Smr’s growth rate is below 0.96. At the left-most point (red box), Em increases in frequency, and the inset plots show that methionine is unlimiting, while arginine is initially limiting. At the right-most point (red box), Em decreases in frequency and the inset plots show that neither methionine nor arginine are limiting at any timepoint. **b**, The rate at which Smr produces arginine also determines coexistence. Em increases in frequency from an initial frequency of 0.1 when the arginine production rate is below 4, but decreases in frequency above this value, indicating that Er takes over the population. Amino acid dynamics at the far left and far right points are similar to the inset plots shown in part a.

Another parameter that we hypothesized could affect amino acid limitation is the rate at which the *E. coli* strains deplete their amino acids (**Fig. S7**). We found that the rate at which Em depletes methionine has no effect on coexistence (**Fig. S7a**), but coexistence is lost when Er’s arginine depletion rate is low (around 25% of the default rate; **Fig. S7b-c**). At a low arginine depletion rate, both amino acids are abundant throughout growth, and Er is able to grow more quickly and outcompete Em. Next, we explored coexistence in a scenario in which the *E. coli* strains deplete both amino acids. In our lab system, it is possible that Em depletes arginine and that Er depletes methionine at low rates. We found that depletion of arginine by Em has no effect (**Fig. S8a**). However, if Er depletes methionine at 90% of the rate at which Em depletes methionine, coexistence is not possible (**Fig. S8b**). If both strains are able to deplete the other strain’s amino acid, the effects cancel out and coexistence is always possible, except when both strains deplete the other amino acid at the same per capita rate at which the auxotroph depletes that amino acid (**Fig. S8c**). We also explored complete overlap in amino acid consumption by creating a model in which both *E. coli* strains require and consume the same amino acid and *S. enterica* only produces this single amino acid. In this situation, the two *E. coli* strains compete for both lactose and the amino acid, and the *E. coli* strains cannot coexist, even if *S. enterica*’s growth rate is low (**Fig. S9**).

Finally, we wondered whether this mechanism promotes coexistence in more complex communities. We added a third *E. coli* amino acid auxotroph (Ef, auxotrophic for phenylalanine) into our model and added production of phenylalanine by *S. enterica* (Smrf) (**Fig. 5a, Table S2**). We set Ef’s growth rate slightly lower than Em’s, and again assessed coexistence by starting each *E. coli* strain rare and tracking whether it could increase in frequency. When Smrf grows more slowly than the *E. coli* strains, all three *E. coli* strains coexist (**Fig. 5b**). However, when Smrf grows faster, the *E. coli* strain with the highest growth rate, Er, outcompetes the other *E. coli* strains (**Fig. 5c**). The mechanism of coexistence is the same as above, where the amino acid consumed by the initially-rare *E. coli* strain(s) is abundant (**Fig. S10**).

**Figure 5.**
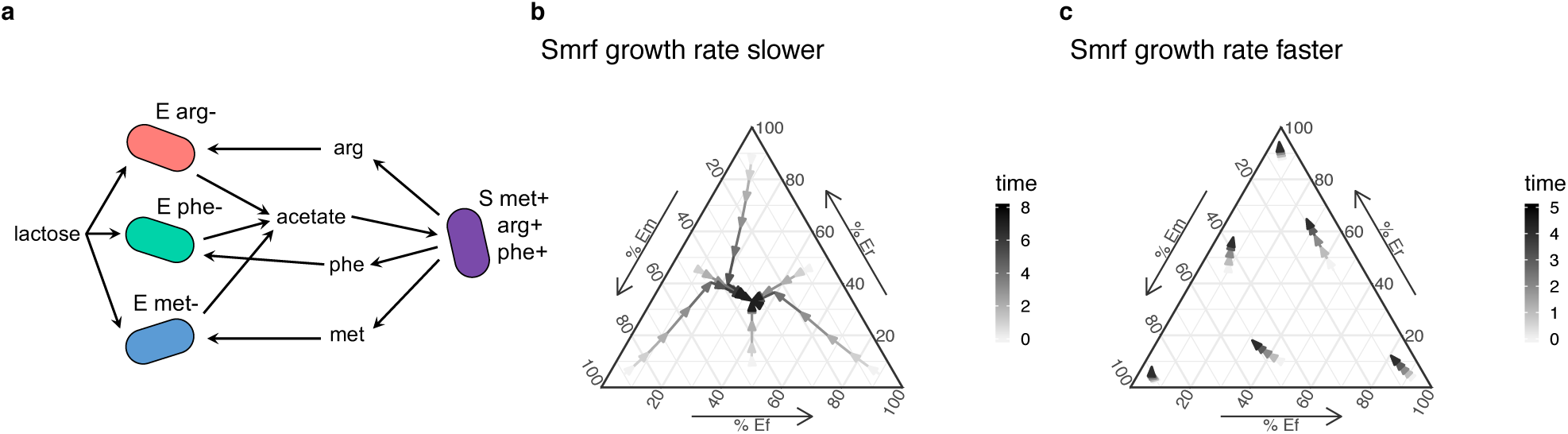
Three competing partner mutualists coexist if the shared mutualist sets the community growth rate. **a**, Schematic showing a community with three *E. coli* partner strains. Ef requires the amino acid phenylalanine, which Smrf supplies in addition to methionine and arginine, which are consumed by Em and Er, respectively. Equations and parameters are described in **Table S2**. The *E. coli* strain growth rates are µ_Em_ = 1, µ_Er_ = 1.1 and µ_Ef_ = 0.9. **b**, A ternary plot showing the frequencies of the three *E. coli* partner strains over time. In these simulations, Smrf’s growth rate is 0.5, lower than all three *E. coli* growth rates, and 10,000 units of lactose were supplied so that larger changes in frequencies could be seen within one growth period. All strains are able to increase in frequency when initially rare, indicating coexistence, because the initially-rare strains’ amino acids are abundant (**Fig. S10**). **c**, When Smrf’s growth rate is 1.5, the frequency of Er increases from all starting frequencies, indicating that Er would take over the population over several growth cycles. 10,000 units of lactose were supplied to show larger changes in frequencies. Growth ceases by timepoint 5, so later timepoints are not shown.

## Discussion

In communities of multiple mutualists, competition between species within a partner guild can affect coexistence and maintenance of diversity^9,10^. We explored the impact of resource competition between partner species that interact with a shared mutualist on coexistence and stability. Laboratory results showed that two *E. coli* partner mutualist strains that receive different amino acids from a *S. enterica* shared mutualist can coexist, despite the fact that one *E. coli* strain is a better competitor for lactose, the resource that ultimately limits growth. Modeling indicated that stability is possible when the *E. coli* strains are temporarily limited by different resources. While lactose sets the total *E. coli* carrying capacity—growth ceases when lactose is exhausted—the availability of amino acids during growth determines the instantaneous growth rate of each *E. coli* strain. When one *E. coli* strain begins rare, its amino acid is always abundant, so its instantaneous growth rate is faster and it consumes lactose quickly. In contrast, the initially-common *E. coli* strain is amino acid-limited at early timepoints and grows more slowly than the rare strain, which allows the initially-rare *E. coli* strain to increase in frequency. The community is therefore stable through negative-frequency dependence. We found that three key parameters affect the potential for coexistence via temporary amino acid limitation. Coexistence is not possible if *S. enterica*’s growth rate is high, if *S. enterica*’s production rate of the initially-common *E. coli*’s amino acid is high, or if the initially-common *E. coli* strain depletes its amino acid at a low rate. In these situations, the initially-common *E. coli* strain is never limited by its amino acid, and the stronger competitor excludes the weaker. In summary, coexistence requires temporary amino acid limitation for one partner strain.

This mechanism of stability is related to classical ideas in ecology about niche partitioning^16,17^. Theoretical work predicts that multiple species are unable to coexist if they are limited by the same resource^11,18^. However, if the species are limited by different resources, they can coexist^19,20^. In our system, the carrying capacity of the *E. coli* strains is ultimately limited by lactose. However, during growth, the limiting resources are temporarily “partitioned.” One strain’s instantaneous growth rate is limited by its amino acid, while the other achieves its maximum growth rate due to an abundance of lactose and its amino acid. An interesting element of our system is that a biotic factor creates the potential for temporary niche partitioning, rather than an aspect of the environment. The shared mutualist, *S. enterica*, causes the two partner species’ instantaneous growth rates to be determined by different resources early on in growth. In addition, the shared mutualist creates the potential for niche partitioning by providing two different resources for the partner strains. Coexistence is not possible if both partner strains receive the same resource from the shared mutualist (**Fig. S9**).

Recent work has explored the importance of competition between partner species in multiple mutualist communities. Several empirical studies have documented competition between species within a partner guild for access to the shared mutualist. For example, flowering plants compete for pollinator services^21^, and multiple species of plant-defending ants compete for nesting sites on host acacia plants^22^. In these cases, partner species may also compete for resources that are external to the mutualism. For example, in the ant-plant systems, plants may compete with one another for water and nutrients, and ants for prey^10^. Johnson and Bronstein (2019) took a mathematical approach to examining the coexistence of two partner mutualists that compete for both a host-provided resource and an external resource^10^. They determined that coexistence requires that one partner is limited by the host-provided resource and the other by the external resource (i.e. niche partitioning). Together with this study, our results suggest that understanding the stability of multiple mutualist systems requires consideration of competition for external resources in addition to competition for access to the shared mutualist. We show that even when both competing partners are ultimately limited by a resource external to the mutualism, coexistence can be maintained through temporary niche partitioning. Our work also identifies the importance of the shared mutualist providing different resources to members of the partner mutualist guild.

Microbial communities are often observed to include many cross-feeding species that exchange metabolites^23–25^. An open question in microbial ecology is why natural communities appear to contain several ecologically-similar species that consume the same resources and carry out the same functions^25,26^. Our work suggests that these communities may be stable despite the potential for competition between strains that provide redundant functions (in our case, the conversion of lactose to acetate). We also showed that temporary limitation by different resources allows for coexistence of three partner strains (**Fig. 5b**), and this mechanism may extend to coexistence of many partner strains. However, there is likely a limit to the number of metabolites that a single shared mutualist can secrete and therefore an upper limit to the system complexity. Other factors that are likely to influence the stability of cross-feeding systems include spatial structure and evolution. In general, spatial structure promotes diversity^27^, though structure can also lead to a loss of strains^28^. Evolution can lead to rapid changes in cross-feeding^29,30^. The evolution of specialists that only interact with a subset of competing partners may decrease the diversity of the system. This will be explored in future work. Finally, our mechanism of coexistence relies on the dynamics created by a batch or seasonal culture regime. However, analytical analysis of a chemostat model of our system indicates that coexistence is also possible in continuous culture, though through a different mechanism (**SI Section 2**).

The results presented here improve our understanding of the ecology of multiple mutualist communities, expanding our knowledge of mutualisms beyond pairwise interactions. Ecological stability is critical for the maintenance of biodiversity. Within mutualistic communities, coexistence of many species within a partner mutualist guild creates functional redundancy, which is important in the face of disturbances because redundancy can protect mutualistic communities from collapse^5,31^. Knowledge of ways to preserve functional redundancy might aid efforts to design stable microbial communities for applications in health and industry^32^.

## Methods

### Strains and media

We used two *Escherichia coli* K12 strains, both derived from the Keio collection^33^. The methionine auxotroph (“Em”) has a Δ*metB* mutation, and the arginine auxotroph (“Er”) has a Δ*argA* mutation. *LacZ* was added to both strains using phage transduction^34^. We also used a *Salmonella enterica* serovar Typhimurium LT2 strain that secretes methionine and arginine (“Smr”). This strain was derived from a strain containing mutations in *metA* and *metJ* that cause over-production of methionine^12,35,36^. We selected for arginine production by coculturing this strain with the *E. coli* arginine auxotroph as a lawn on lactose minimal media plates containing x-gal (0.05% v/v) for four 7-day growth cycles with 1:6.67 dilutions at each transfer^37^. The appearance of a blue colony suggested the evolution of arginine production in *S. enterica*, which we confirmed by isolating *S. enterica* from the colony on citrate minimal media plates and cross-streaking with Er. We sequenced this strain using Illumina NextSeq and identified mutations using breseq^38^. We found a T→A point mutation in *argG* at position 3459818 (reference strain NC_003197).

In coculture or three-strain cultures, strains were grown in a modified Hypho minimal medium with lactose as the carbon source, containing 2.78 mM lactose, 14.5 mM K_2_HPO_4_, 16.3 mM NaH_2_PO_4_, 0.814 mM MgSO4, 3.78 mM Na_2_SO_4_, 3.78 mM [NH_4_]_2_SO_4_, and trace metals (1.2 µM ZnSO_4_, 1 µM MnCl_2_, 18 µM FeSO_4_, 2 µM (NH_4_)_6_Mo_7_O_24_, 1 uM CuSO_4_, 2 mM CoCl_2_, 0.33 µm Na_2_WO_4_, 20 µM CaCl_2_). Monocultures and cocultures of Em and Er were grown in this medium with 250 µM of methionine and 250 µM of arginine added, concentrations that we found to be unlimiting (i.e. growth ceased when lactose was depleted, rather than the amino acids; **Fig. S1**). Smr’s growth rate in monoculture was assessed in Hypho minimal medium with 12 mM acetate rather than lactose, a concentration that approximates the total amount of acetate produced by the *E. coli* strains.

To measure final yields as colony-forming units, cultures were diluted in saline (0.85% NaCl) and plated on Hypho minimal media plates with 1% agar. Plates for the *E. coli* strains contained 2.78 mM lactose and 100 µM of methionine for Em, or 100 µM of arginine for Er. Smr was plated on Hypho plates containing 3.4 mM sodium citrate instead of lactose. All plates contained 0.05% v/v x-gal, which makes *E. coli* colonies blue.

### Growth assays

All experiments were performed in 96-well plates with 200 µl of media per well, inoculated with a 1:200 dilution of log-phase monocultures (1 µl of each strain). We measured OD600 in a Tecan InfinitePro 200 plate reader at 30°C, shaking at 432 rpm between readings, which were taken every 20 minutes. Growth rate estimates were calculated by fitting growth curves to a Baranyi function^39^ by obtaining nonlinear least-square estimates and using the growth rate parameter estimate.

### Mutual invasibility experiments

The ability of each *E. coli* strain to increase in frequency from rare was our criterion for coexistence^13,14^. For these experiments, initial Smr density was kept constant (9.8e5 CFU/ml), and the *E. coli* total density was kept constant (1.6e6 CFU/ml) but the frequency of each strain differed across a range of three frequencies—0.024, 0.320, and 0.899 Em/(Em + Er). After growth, the cultures were diluted and plated to measure yields as CFU/ml (media described above). The change in Em frequency was calculated as [Em/ (Em + Er)]_final_ – [Em/(Er + Er)]_initial_.

### Ecological modeling

The ecological model is shown in Results and **Table S1**. The ODE system was solved using the deSolve package in R, which used the lsoda solver to numerically integrate. All simulations were solved for sufficient duration to ensure dynamics had ceased. During integration, relative tolerance (rtol) was set to 1e-13 and maxsteps to 1e5.

## Analysis & Statistics

Modeling, data visualization, and statistical analyses were done in R version 3.6.0. Growth rates were compared using a one-way analysis of variance (ANOVA) and post hoc Tukey HSD tests, and other tests were two-sided t-tests, all with α = 0.05. We used the R package ggtern to make the ternary plots in **Fig. 5**.

## Acknowledgements

The authors thank Lisa Fazzino, Ben Holte, and Alejandro Behling for assistance with preliminary experiments, Jeremy Chacón for help with modeling, and Brian Smith, Leno Smith, and Jonathan Martinson for providing comments on the manuscript. This work was supported by a National Science Foundation Graduate Research Fellowship for S.P.H. and by the National Institutes of Health (1R01-GM121498, to W.R.H.).

## Data Availability

The data that support the findings of this study are available from the corresponding author upon request.

## Author Contributions

S.P.H. conceived the study, collected and analyzed the data, and prepared the manuscript. W.H. helped design the experiments, oversaw data analysis, supplied materials, and assisted with manuscript preparation. T.G. analyzed the chemostat model, and provided comments on the manuscript. R.C. constructed the Er strain, provided ideas for additional modeling experiments, and provided comments on the manuscript.

