## Supplementary Information for "Limitation by a shared mutualist promotes coexistence of multiple competing partners"

### Supplementary Information for Hammarlund et al.

#### Contents

Section 1: Supplementary Figures 1-10, Tables 1 & 2

Section 2: Chemostat Analysis

#### Section 1

**Figure 1 | Amino acid requirements of Em and Er.** Final yields (OD600) of **a**, Em across a range of methionine concentrations and **b**, Er monocultures across a range of arginine concentrations, in lactose minimal media. Cultures were inoculated with log-phase culture and grown until stationary phase was reached, then OD was measured (see Methods for growth conditions and OD measurements). Yields plateau at 100 µM of methionine for Em and between 100 µM and 500 µM of arginine for Er, suggesting that Em requires slightly less methionine than Er requires arginine to reach its maximum yield. We chose 250µM as the concentration of both amino acids for our lactose minimal media supplemented with arginine and methionine because yields saturate around this concentration, indicating that lactose sets carrying capacity rather than the amino acid.

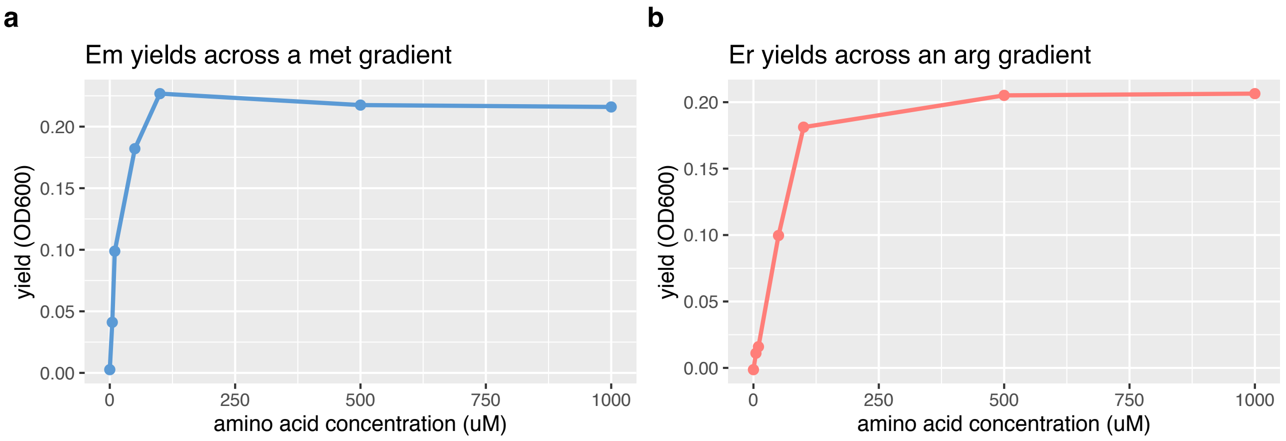

**Figure 2 | Monoculture final yields (CFU/ml).** Yields of Em, Er, and Smr in monoculture, measured by diluting and plating on selective media. Em and Er were grown in lactose media with methionine and arginine, and Smr in acetate media (see Methods).

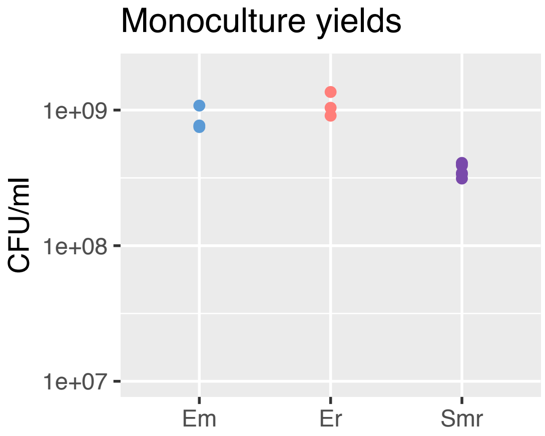

**Figure 3 | Coculture final yields (CFU/ml). a**, Yields of Em and Smr from Em+Smr cocultures, grow in lactose minimal media, and measured by diluting and plating on selective media. **b**, Yields of Er and Smr from Er+Smr cocultures, also grown in lactose media and diluted and plated to count colonies.

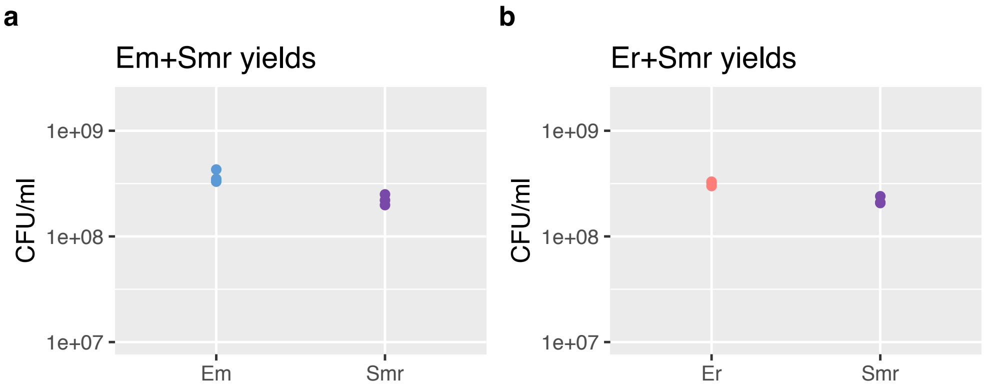

**Figure 4 | Yields (CFU/ml) of Smr Em Er communities across a range of initial Em frequencies.** Cultures were diluted and plated onto selective media, and colonies were counted. **a**, Em yields (data shown as Em frequencies in Fig. 1e). **b**, Er yields (data shown as Em frequencies in Fig. 1e). **c**, Smr yields.

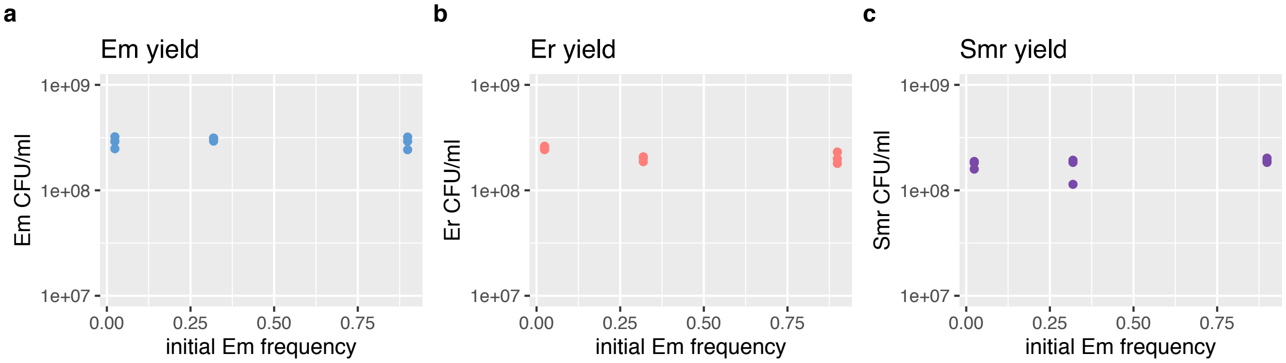

**Table 1 | Model parameters and default values.** Explanation of parameter names, units, defaults, and biological interpretations.

| **Parameter or state variable** | **Units** | **Default or initial value[1]** | **Name** | **Biological Interpretation** |
|---|---|---|---|---|
| Smr | Cells/ml | 100 at start | Smr cell density | *S. enterica* methionine and arginine producer population size |
| Em | Cells/ml | variable at start | Em cell density | *E. coli* methionine auxotroph population size |
| Er | Cells/ml | variable at start | Er cell density | *E. coli* arginine auxotroph population size |
| lcts | Cells/ml | 1000 at start | lactose | The concentration of lactose that can produce a certain cell density once consumed (i.e. cell equivalents)[2] |
| met | Cells/ml | 1 at start | methionine | The concentration of methionine that can produce a certain cell density once consumed (i.e. cell equivalents)[2] |
| arg | Cells/ml | 1 at start | arginine | The concentration of arginine that can produce a certain cell density once consumed (i.e. cell equivalents)[2] |
| ac | Cells/ml | 1 at start | acetate | The concentration of acetate that can produce a certain cell density once consumed (i.e. cell equivalents)[2] |
| $\mu_{Smr}$ | 1/Timestep | 0.5 | Smr growth rate | Maximum growth rate of Smr |
| $\mu_{Em}$ | 1/Timestep | 1.0 | Em growth rate | Maximum growth rate of Em |
| $\mu_{Er}$ | 1/Timestep | 1.1 | Er growth rate | Maximum growth rate of Er |
| $k_{Em\ lcts}$<br>$k_{Er\ lcts}$<br>$k_{Em\ met}$<br>$k_{Er\ arg}$<br>$k_{Smr\ ac}$ | Cells/ml | 0.01 | Monod half-saturation constants | The value of k determines the resource concentration where growth rate is half-maximum |
| $p_{Smr\ met}$<br>$p_{Smr\ arg}$<br>$p_{Em\ ac}$<br>$p_{Er\ ac}$ | Cells/cell | 1.0001 | Resource production rates | The amount of the specified resource produced by the specified strain, in cell equivalents per cell |

| | | | | |
|---|---|---|---|---|
| $c_{Em\ met}$ [3]<br>$c_{Er\ arg}$ [3] | Unitless | 1 | Amino acid depletion rate | A term that scales the depletion of methionine by Em and arginine by Er, relative to a baseline depletion of 1 |
| $c_{Em\ arg}$ [4]<br>$c_{Er\ met}$ [4] | Unitless | 0 | Depletion rate of the other *E. coli* strain's required amino acid | A term that scales the depletion of arginine by Em and methionine by Er, relative to a baseline depletion of 0 |

[1] Analyses with altered parameter values are stated in the results and figure legends.

[2] Resources are defined in terms of the amount of cells that can be produced by that concentration ("cell equivalents"). Specifically, resource concentrations (mmol resource/ml) are multiplied by a conversion parameter (1 cell/mmol resource) to result in cells/ml. For example, 1000 cell equivalents of lactose means that *E. coli* would grow to a density of 1000 cells/ml.

[3] Amino acid equations using these depletion rates are shown below with Supplementary Figure 7.

[4] Amino acid equations using these depletion rates are shown below with Supplementary Figure 8.

**Figure 5 | Dynamics of strains and all resources when Em starts rare.** Growth curves and resource plots at the point labeled “**c**” in Fig. 3b.

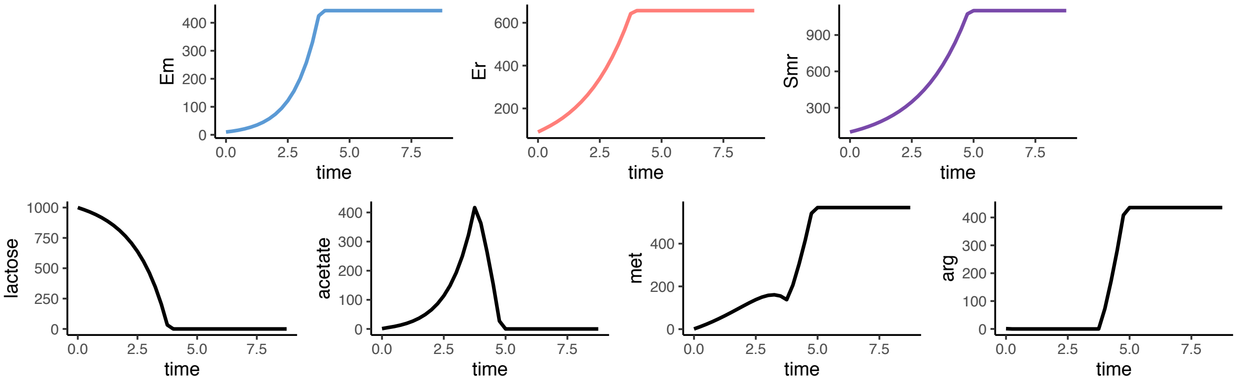

**Figure 6 | Dynamics of strains and all resources when Em starts rare.** Growth curves and resource plots at the point labeled "**d**" in Fig. 3b.

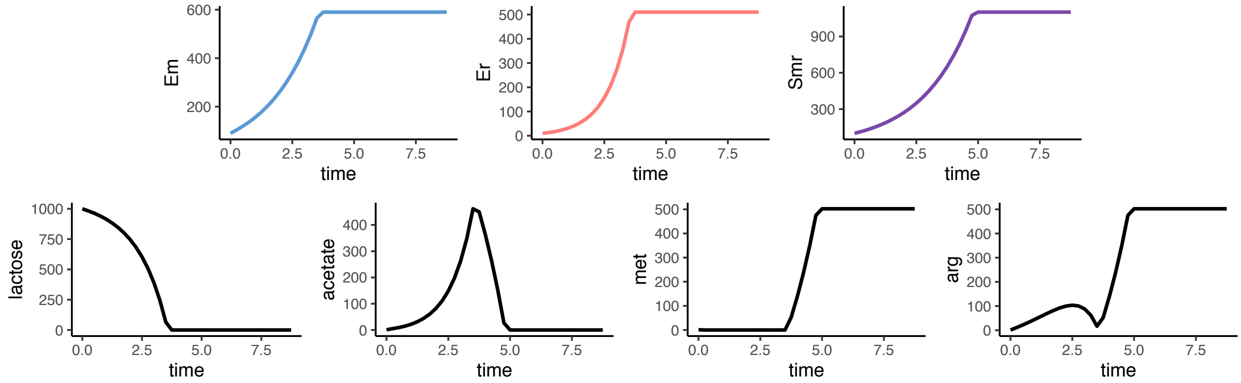

**Figure 7 | Coexistence is robust to changes in amino acid depletion rates.**
Depletion rates $c_{Em\ met}$ and $c_{Er\ arg}$ adjust the rate at which the *E. coli* strains deplete their required amino acids. The default value is 1. A change in the depletion rate does not directly affect the *E. coli* strains' growth (i.e. the equations for Em and Er are unchanged). See Supplementary Table 1 for further description of these parameters and the equations below for how they are incorporated into the model. **a**, Effects of altering the met depletion rate by Em only. Em is always able to invade from rare. **b**, Effects of altering the arg depletion rate by Er only. Coexistence is maintained except at values of $c_{Er\ arg}$ below 0.26. **c**, The effects of altering both depletion rates, keeping them equal. Coexistence is possible above $c_{Em\ met} = c_{Er\ arg} = 0.26$. For all plots, the dotted vertical line at x = 1 shows default value of the depletion rate.

Altered equations for amino acids using these depletion rates (in bold):

$$\frac{d\ met}{dt} = -\left(\frac{dEm}{dt} \times \boldsymbol{c_{Em\ met}}\right) + \left(p_{Smr\ met} \times \frac{dSmr}{dt}\right)$$

$$\frac{d\ arg}{dt} = -\left(\frac{dEr}{dt} \times \boldsymbol{c_{Er\ arg}}\right) + \left(p_{Smr\ arg} \times \frac{dSmr}{dt}\right)$$

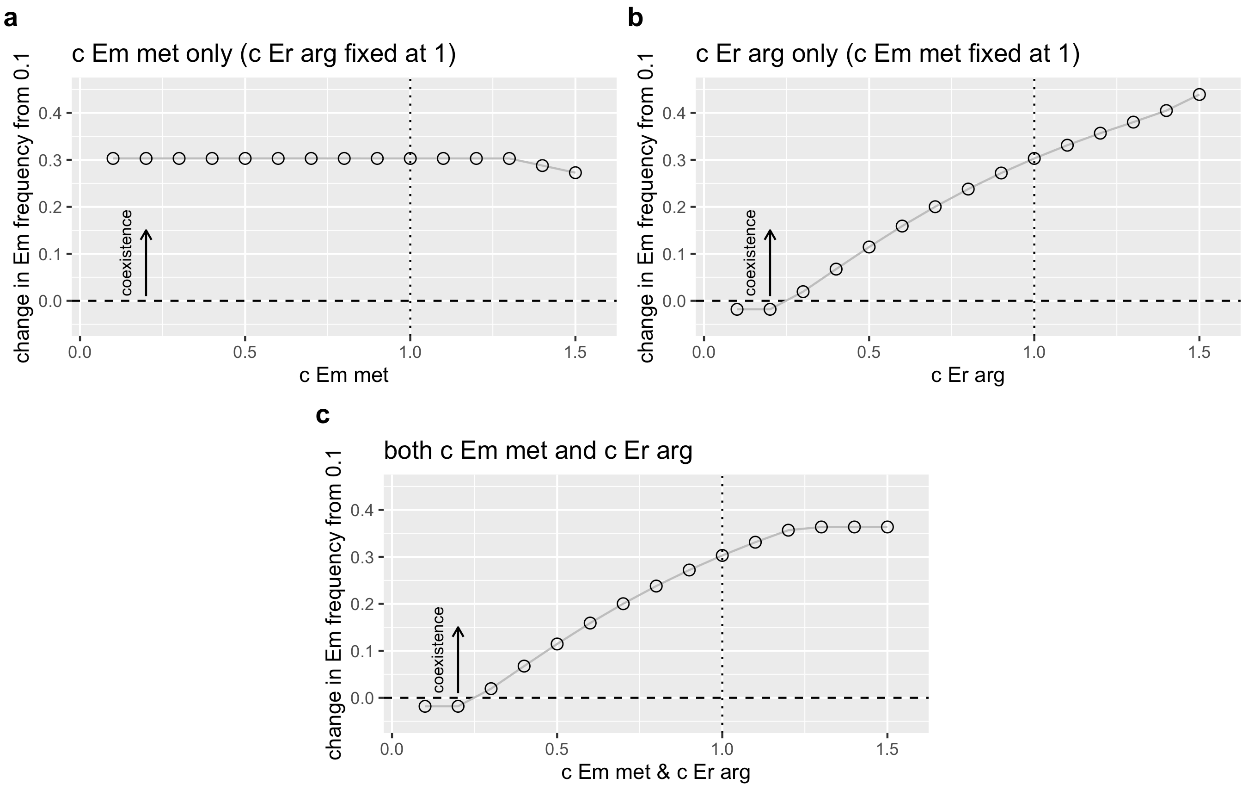

**Figure 8 | Coexistence is robust to depletion of both amino acids by both *E. coli* strains.** Depletion rates $c_{Em\ arg}$ and $c_{Er\ met}$ range between 0 and 1 and adjust the rate at which each *E. coli* strain depletes the other *E. coli* strain's required amino acid. The non-required amino acid does not contribute to the population density of the strain (i.e. the equations for Em and Er are unchanged), which is supported by lab data. The default values of $c_{Em\ arg}$ and $c_{Er\ met}$ are 0. See Supplementary Table 1 for further description of these parameters and the equations below for how they are incorporated into the model. **a**, The effect of $c_{Em\ arg}$, while $c_{Er\ met}$ is held at its default value of 0. Coexistence is maintained across the whole range from 0 to 1. **b**, The effect of $c_{Er\ met}$, while $c_{Em\ arg}$ is held at its default value of 0. Coexistence is lost at $c_{Er\ met} = 0.9$ and above. **c**, Both strains deplete the other strain's amino acid at the same rate. Between 0 and 0.9, the effects of $c_{Em\ arg}$ and $c_{Er\ met}$ cancel one another out and coexistence is maintained, but at $c_{Em\ arg} = c_{Er\ met} = 1$, coexistence is lost. For all plots, the dotted vertical line at x = 0 shows the default value of the depletion rate.

Altered equations for amino acids using these depletion rates (in bold):

$$\frac{d\ met}{dt} = -\frac{dEm}{dt} + \left(p_{Smr\ met} \times \frac{dSmr}{dt}\right) - \left(\boldsymbol{c_{Er\ met}} \times \frac{dEr}{dt}\right)$$

$$\frac{d\ arg}{dt} = -\frac{dEr}{dt} + \left(p_{Smr\ arg} \times \frac{dSmr}{dt}\right) - \left(\boldsymbol{c_{Em\ arg}} \times \frac{dEm}{dt}\right)$$

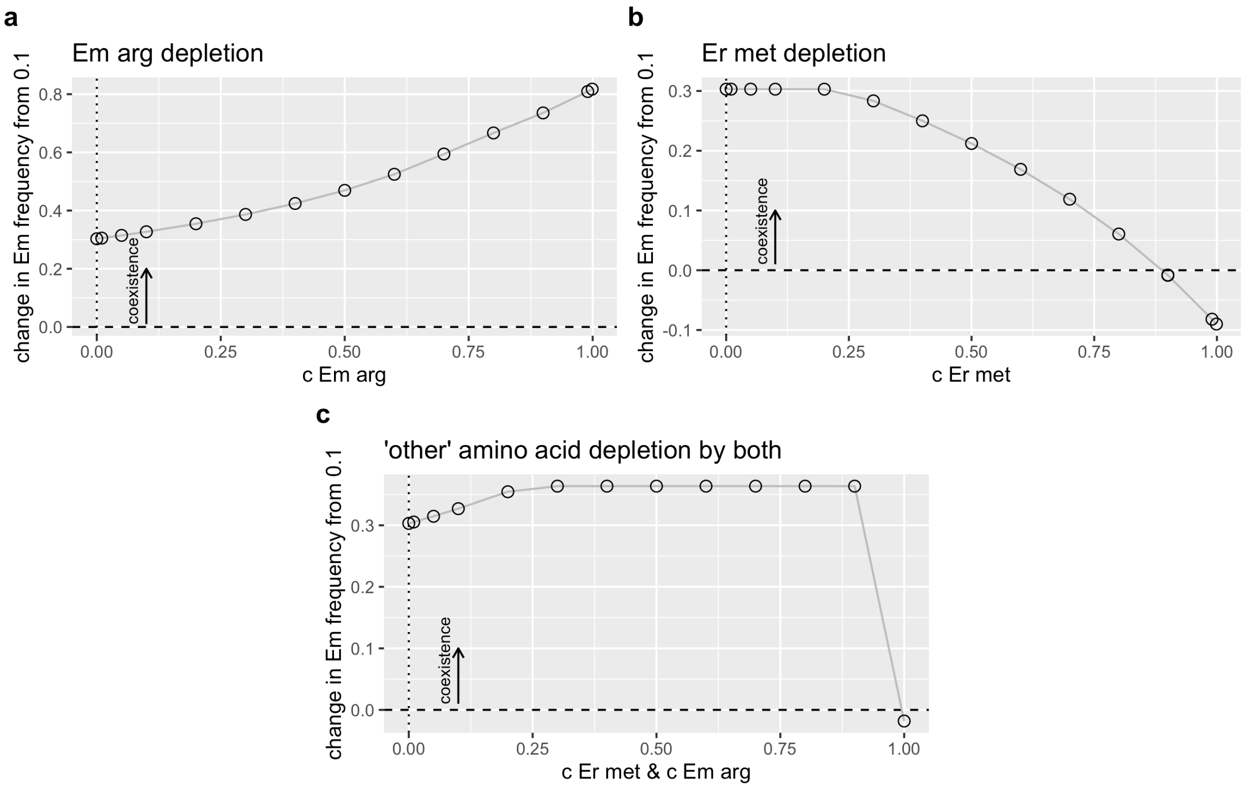

**Figure 9 | If *S. enterica* provides methionine to two different methionine-requiring *E. coli* strains (Em1, Em2) there is no coexistence, even if Sm grows more slowly. a**, Schematic of the community. E met-1 (Em1) and E met- 2 (Em2) both require methionine. S met+ (Sm) produces methionine. The *E. coli* strains produce acetate, which Sm consumes, and both *E. coli* strains consume lactose. The equations are shown below. **b**, When Sm grows more slowly ($\mu_{Em1} = 1$, $\mu_{Em2} = 1.1$, $\mu_{Sm} = 0.5$), there is no coexistence. The slower growing Em strain (Em1) cannot increase in frequency from any starting frequency. **c**, The same is true when Sm grows faster than the two Em strains ($\mu_{Em1} = 1$, $\mu_{Em2} = 1.1$, $\mu_{Sm} = 1.5$). Em1 decreases in frequency from every starting frequency. **d**, Timeseries showing met limitation at early timepoints where $\mu_{Em1} = 1$, $\mu_{Em2} = 1.1$, $\mu_{Sm} = 0.5$ and the initial frequency of Em1 is 0.1. The two *E. coli* strains compete for both methionine and lactose—methionine limits instantaneous growth rates until lactose is depleted and growth ceases.

The equations are similar to the three-strain model, except there are two *E. coli* strains that consume methionine, Em1 and Em2. Parameters are as described in Supplementary Table 1.

$$\frac{d\,Em1}{dt} = Em1 \times \mu_{Em1} \times \frac{lcts}{lcts + k_{Em1\,lcts}} \times \frac{met}{met + k_{Em1\,met}}$$

$$\frac{d\,Em2}{dt} = Em2 \times \mu_{Em2} \times \frac{lcts}{lcts + k_{Em2\,lcts}} \times \frac{met}{met + k_{Em2\,met}}$$

$$\frac{d\,Sm}{dt} = Sm \times \mu_{Sm} \times \frac{ac}{ac + k_{Sm\,ac}}$$

$$\frac{d\,lcts}{dt} = -\frac{dEm1}{dt} - \frac{dEm2}{dt}$$

$$\frac{d\,ac}{dt} = \left(p_{Em1\,ac} \times \frac{dEm1}{dt}\right) + \left(p_{Em2\,ac} \times \frac{dEm2}{dt}\right) - \frac{dSm}{dt}$$

$$\frac{d\,met}{dt} = -\frac{dEm1}{dt} - \frac{dEm2}{dt} + \left(p_{Sm\,met} \times \frac{dSm}{dt}\right)$$

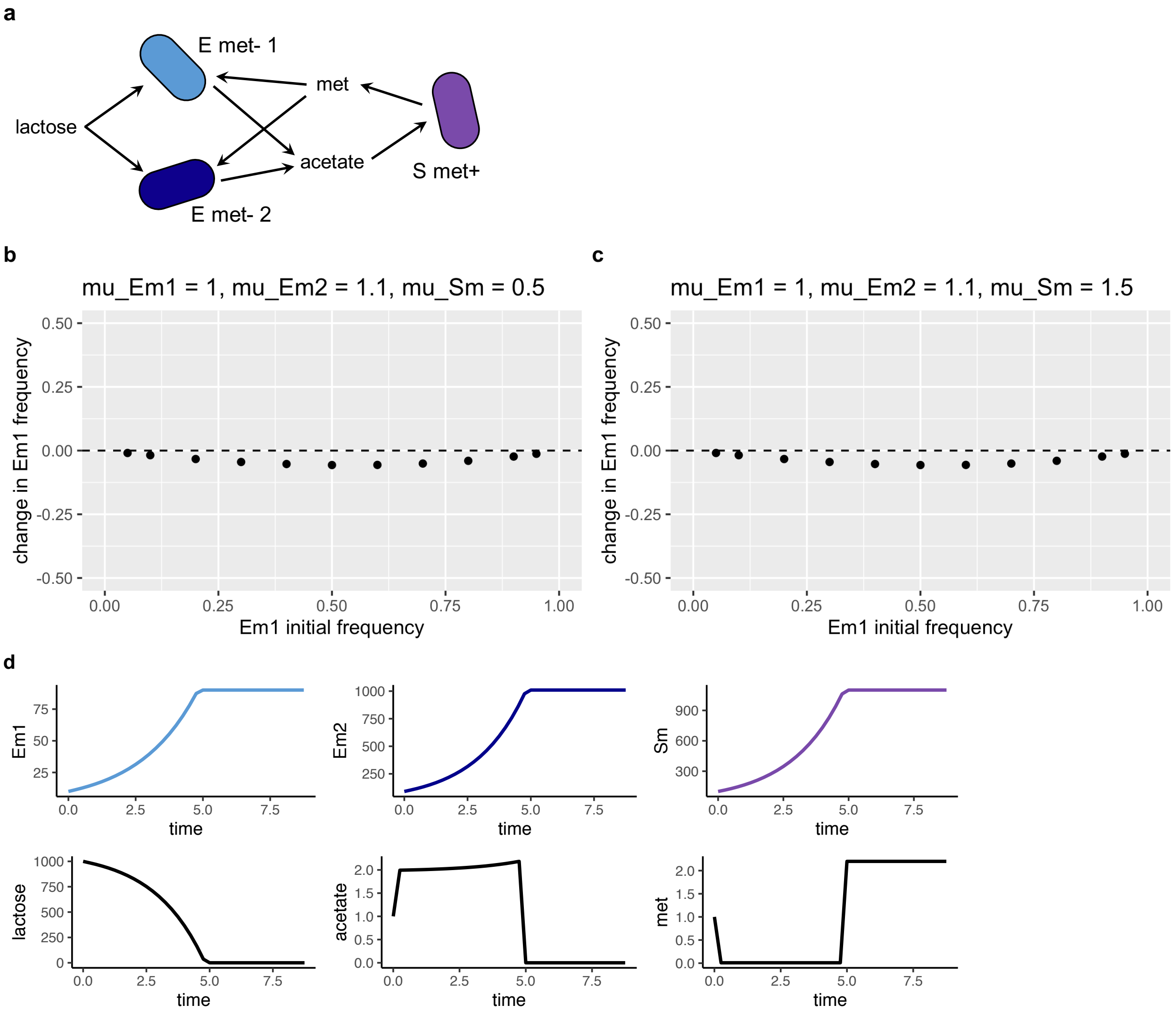
a
E met- 1
met
lactose
acetate
S met+
E met- 2
b
mu_Em1 = 1, mu_Em2 = 1.1, mu_Sm = 0.5
change in Em1 frequency
Em1 initial frequency
c
mu_Em1 = 1, mu_Em2 = 1.1, mu_Sm = 1.5
change in Em1 frequency
Em1 initial frequency
d
Em1
Em2
Sm
lactose
acetate
met
time

**Table 2 | Model parameters and default values for three *E. coli* strain community.** Explanation of parameter names, units, defaults, and biological interpretations. Equations are shown below the table.

| Parameter or state variable | Units | Default or initial value[1] | Name | Biological Interpretation |
|---|---|---|---|---|
| Smrf | Cells/ml | 100 at start | Smr cell density | *S. enterica* methionine, arginine, and phenylalanine producer population size |
| Em | Cells/ml | variable at start | Em cell density | *E. coli* methionine auxotroph population size |
| Er | Cells/ml | variable at start | Er cell density | *E. coli* arginine auxotroph population size |
| Ef | Cells/ml | variable at start | Ef cell density | *E. coli* phenylalanine auxotroph population size |
| lcts | Cells/ml | 1000 at start | lactose | The concentration of lactose that can produce a certain cell density once consumed (i.e. cell equivalents)[2] |
| met | Cells/ml | 1 at start | methionine | The concentration of methionine that can produce a certain cell density once consumed (i.e. cell equivalents)[2] |
| arg | Cells/ml | 1 at start | arginine | The concentration of arginine that can produce a certain cell density once consumed (i.e. cell equivalents)[2] |
| phe | Cells/ml | 1 at start | phenylalanine | The concentration of phenylalanine that can produce a certain cell density once consumed (i.e. cell equivalents)[2] |
| ac | Cells/ml | 1 at start | acetate | The concentration of acetate that can produce a certain cell density once consumed (i.e. cell equivalents)[2] |
| $\mu_{Smrf}$ | 1/Timestep | 0.5 | Smr growth rate | Maximum growth rate of Smr |
| $\mu_{Em}$ | 1/Timestep | 1.0 | Em growth rate | Maximum growth rate of Em |
| $\mu_{Er}$ | 1/Timestep | 1.1 | Er growth rate | Maximum growth rate of Er |
| $\mu_{Ef}$ | 1/Timestep | 0.9 | Ef growth rate | Maximum growth rate of Ef |
| $k_{Em\ lcts}$<br>$k_{Er\ lcts}$<br>$k_{Ef\ lcts}$ | Cells/ml | 0.01 | Monod half-saturation constants | The value of k determines the resource concentration where growth rate is half-maximum |

| | | | | |
|---|---|---|---|---|
| $k_{Em\ met}$<br>$k_{Er\ arg}$<br>$k_{Ef\ phe}$<br>$k_{Smrf\ ac}$ | | | | |
| $p_{Smrf\ met}$<br>$p_{Smrf\ arg}$<br>$p_{Smrf\ phe}$<br>$p_{Em\ ac}$<br>$p_{Er\ ac}$<br>$p_{Ef\ ac}$ | Cells/cell | 1.0001 | Resource production rates | The amount of the specified resource produced by the specified strain, in cell equivalents per cell |

[1] Analyses with altered parameter values are stated in the results and figure legends.

[2] Resources are defined in terms of the amount of cells that can be produced by that concentration ("cell equivalents"). Specifically, resource concentrations (mmol resource/ml) are multiplied by a conversion parameter (1 cell/mmol resource) to result in cells/ml. For example, 1000 cell equivalents of lactose means that *E. coli* would grow to a density of 1000 cells/ml.

Three *E. coli* strain model equations:

$$\frac{d\,Em}{dt} = Em \times \mu_{Em} \times \frac{lcts}{lcts + k_{Em\ lcts}} \times \frac{met}{met + k_{Em\ met}}$$

$$\frac{d\,Er}{dt} = Er \times \mu_{Er} \times \frac{lcts}{lcts + k_{Er\ lcts}} \times \frac{arg}{arg + k_{Er\ arg}}$$

$$\frac{d\,Ef}{dt} = Ef \times \mu_{Ef} \times \frac{lcts}{lcts + k_{Ef\ lcts}} \times \frac{phe}{phe + k_{Ef\ phe}}$$

$$\frac{d\,Smrf}{dt} = Smrf \times \mu_{Smrf} \times \frac{ac}{ac + k_{Smrf\ ac}}$$

$$\frac{d\,lcts}{dt} = -\frac{dEm}{dt} - \frac{dEr}{dt} - \frac{dEf}{dt}$$

$$\frac{d\,ac}{dt} = \left(p_{Em\ ac} \times \frac{dEm}{dt}\right) + \left(p_{Er\ ac} \times \frac{dEr}{dt}\right) + \left(p_{Ef\ ac} \times \frac{dEf}{dt}\right) - \frac{dSmrf}{dt}$$

$$\frac{d\,met}{dt} = -\frac{dEm}{dt} + \left(p_{Smrf\ met} \times \frac{dSmrf}{dt}\right)$$

$$\frac{d\ arg}{dt} = -\frac{dEr}{dt} + \left(p_{Smrf\ arg} \times \frac{dSmrf}{dt}\right)$$

$$\frac{d\ phe}{dt} = -\frac{dEf}{dt} + \left(p_{Smrf\ phe} \times \frac{dSmrf}{dt}\right)$$

**Figure 10 | In the three *E. coli* strain community, the rare *E. coli* strain's amino acid is abundant.** Amino acid dynamics are shown for three different initial *E. coli* strain frequencies. **a**, Em is initially rare (at start, Em = 10, Er = 45, Ef = 45) and methionine is abundant throughout growth. **b**, Er is initially rare (at start, Em = 45, Er = 10, Ef = 45) and arginine is abundant throughout growth. **c**, Ef is initially rare (at start, Em = 45, Er = 45, Ef = 10) and phenylalanine is abundant throughout growth. **d**, When two strains begin rare (at start, Em = 5, Er = 5, Ef = 90), both initially-rare strains' amino acids (methionine and phenylalanine) are abundant throughout growth. The two other combinations of rare strains are not shown because dynamics are identical—the rare strains' amino acids are abundant and the common strain's amino acid is limiting.

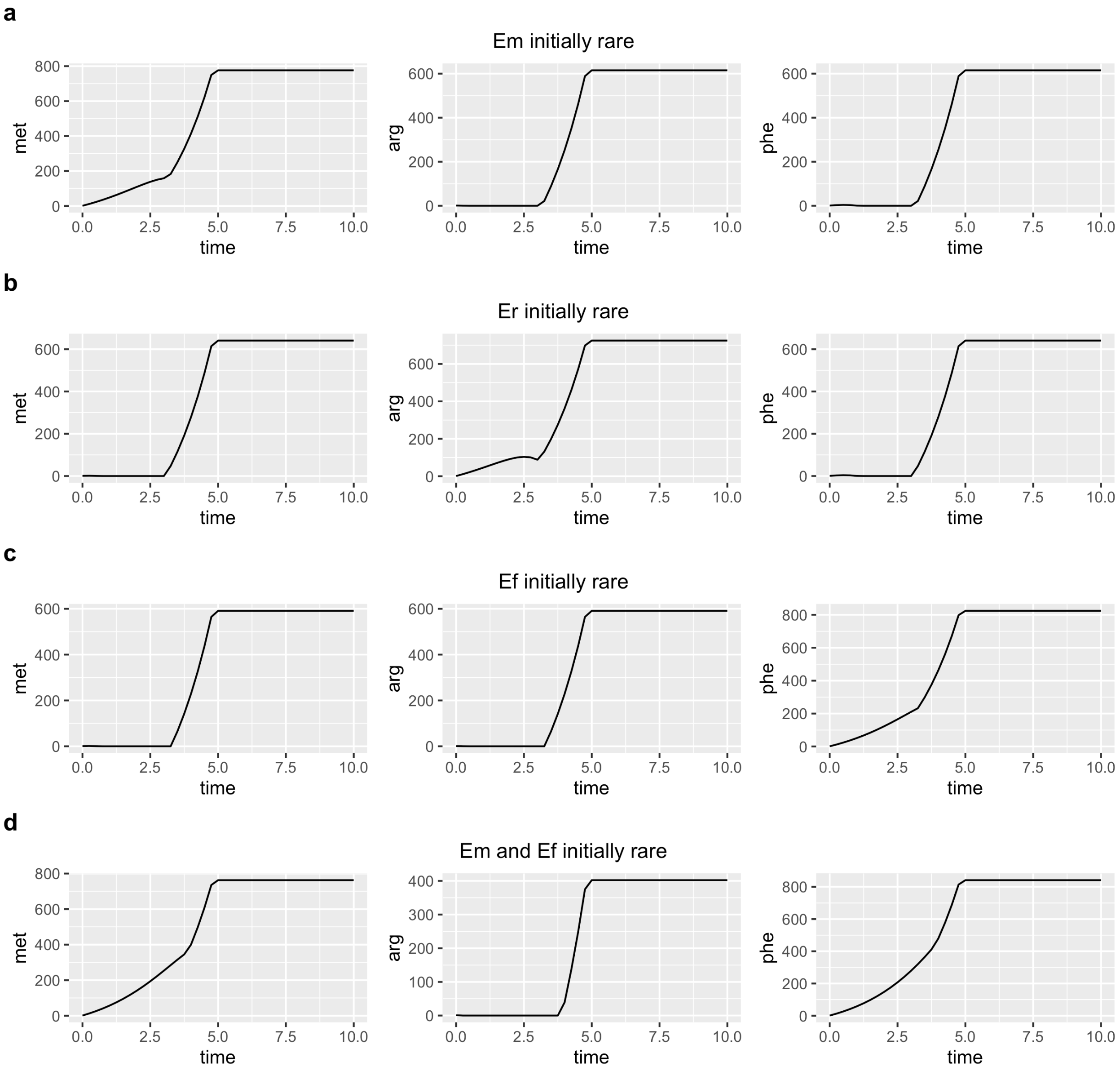

#### Section 2

### COEXISTENCE IN A CHEMOSTAT

#### 1. Chemostat Model

We consider an analogous model in a chemostat with an influx rate $L$ of lactose, and dilution rate $\gamma$.

We first summarize the results.

- There is always a stable steady state that corresponds to the washout of all microbial strains from the chemostat.
- There are steady states of one *E. coli* strain coexisting with *S. enterica*. These states require that the growth rates of each strain are high enough to match the dilution rate of the chemostat.
- The steady state with the slower *E. coli* and *S. enterica* can always be invaded by an infinitesimal concentration of the *E. coli* strain with a faster growth rate.
- The steady state of the faster *E. coli* with *S. enterica* can be invaded by the slower *E. coli* when the amino acid necessary for the slower grower is in sufficient abundance to overcome the advantage of the faster grower.
- The coexistence steady state for all three community members, when it exists, induces bistability in the phase space. The inoculum size determines whether populations die out, or converge to the stable coexistence steady state. There is also an unstable steady state whose stable manifold divides the basins of attraction of coexistence and washout steady states.
- The approach to the steady state may involve multiple dramatic oscillations in the community composition.

We define functions

$$f_{Em}(lcts) = \frac{lcts}{lcts + k_{Em,lcts}},$$
$$f_{Er}(lcts) = \frac{lcts}{lcts + k_{Er,lcts}},$$
$$g(met) = \frac{met}{met + k_{Em,met}},$$
$$h(arg) = \frac{arg}{arg + k_{Er,arg}},$$
$$F(ac) = \frac{ac}{ac + k_{Smr,ac}}$$

and consider the following model

$$\begin{aligned}
\dot{E}_m &= E_m(\mu_{Em} f_{Em}(lcts)g(met) - \gamma)\\
\dot{E}_r &= E_r(\mu_{Er} f_{Er}(lcts)h(arg) - \gamma)\\
\dot{Smr} &= Smr(\mu_{Smr} F(ac) - \gamma)\\
\dot{lcts} &= -E_m\mu_{Em} f_{Em}(lcts)g(met) - E_r\mu_{Er} f_{Er}(lcts)h(arg) - \gamma lcts + L\\
\dot{ac} &= p_{Em,ac}E_m\mu_{Em} f_{Em}(lcts)g(met) + p_{Er,ac}E_r\mu_{Er} f_{Er}(lcts)h(arg) - Smr\mu_{Smr}F(ac) - \gamma ac\\
\dot{met} &= -\mu_{Em}E_m f_{Em}(lcts)g(met) + p_{Smr,met}\mu_{Smr}SmrF(ac) - \gamma met\\
\dot{arg} &= -\mu_{Er}E_r f_{Er}(lcts)h(arg) + p_{Smr,arg}\mu_{Smr}SmrF(ac) - \gamma arg.
\end{aligned}$$

#### 2. Parameters and assumptions

We are interested in finding steady states for this system.

In order to facilitate our analysis, we simplify the notation using parameter choices that were used to simulate batch experiments. Note however that the growth rates for the two *E. coli* strains are flipped compared to the main text. Here, the methionine auxotroph grows faster. The identity of the faster growing strain is arbitrary, and the opposite assumption will lead to analogous results with the *E. coli* strains' identities exchanged.

$$\begin{aligned}
k_{Em,lcts} &= k_{Er,lcts} = k_{Em,met} = k_{Em,arg} = k_{Smr,ac} = 0.01\\
p_{Em,ac} &= p_{Er,ac} = p_{Smr,met} = p_{Smr,arg} = 1.0001\\
\mu_{Smr} &= 0.5;\\
\mu_{Er} &= 1;\\
\mu_{Em} &= 1.1.
\end{aligned} \tag{1}$$

In particular we will make the following assumptions:

**(a):** Since the half-saturation constants are the same, all the Hill functions in the problem are identical, only evaluated at different substrates. Therefore we write

$$f(x) := f_{Em}(x) = f_{Er}(x) = g(x) = h(x) = F(x).$$

This assumption is not essential to our arguments about the existence of various steady states, as our arguments only depend on the monotonicity of the Hill functions.

**(b):** We assume that production rates of amino acids by *S. enterica* are the same

$$p := p_{Smr,arg} = p_{Smr,met}.$$

**(c):** We assume that $\mu_{Em} > \mu_{Er}$. The opposite assumption will lead to analogous results with the *E. coli* strains' identities reversed.

Further, we use a shortened version of the variables above and write $S$ for $Smr$, $\ell$ for $lcts$, $c$ for $ac$, $m$ for $met$ and $a$ for $arg$. Then the system becomes

$$\begin{aligned}
\dot{E}_m &= E_m(\mu_{Em} f(\ell) f(m) - \gamma) \\
\dot{E}_r &= E_r(\mu_{Er} f(\ell) f(a) - \gamma) \\
\dot{S} &= S(\mu_{Smr} f(c) - \gamma) \\
\dot{\ell} &= -E_m \mu_{Em} f(\ell) f(m) - E_r \mu_{Er} f(\ell) f(a) - \gamma\ell + L \\
\dot{c} &= p_{Em,ac} E_m \mu_{Em} f(\ell) f(m) + p_{Er,ac} E_r \mu_{Er} f(\ell) f(a) - S \mu_{Smr} f(c) - \gamma c \\
\dot{m} &= -\mu_{Em} E_m f(\ell) f(m) + p \mu_{Smr} S f(c) - \gamma m \\
\dot{a} &= -\mu_{Er} E_r f(\ell) f(a) + p \mu_{Smr} S f(c) - \gamma a.
\end{aligned} \tag{2}$$

#### 3. Steady states

In this section, we compute the steady states of the chemostat. Our analysis will focus on the existence of different steady states and their stability. Where we can, we will make general statements valid for all parameters; where we cannot make general statements, we will provide formulas in terms of parameters. Our last resort will be to provide numerical support for our assertions for the parameters that we use for the model.

The steady state equations are

$$\begin{aligned}
0 &= E_m(\mu_{Em} f(\ell) f(m) - \gamma) \\
0 &= E_r(\mu_{Er} f(\ell) f(a) - \gamma) \\
0 &= S(\mu_{Smr} f(c) - \gamma) \\
0 &= -E_m \mu_{Em} f(\ell) f(m) - E_r \mu_{Er} f(\ell) f(a) - \gamma\ell + L \\
0 &= p_{Em,ac} E_m \mu_{Em} f(\ell) f(m) + p_{Er,ac} E_r \mu_{Er} f(\ell) f(a) - S \mu_{Smr} f(c) - \gamma c \\
0 &= -\mu_{Em} E_m f(\ell) f(m) + p \mu_{Smr} S f(c) - \gamma m \\
0 &= -\mu_{Er} E_r f(\ell) f(a) + p \mu_{Smr} S f(c) - \gamma a.
\end{aligned} \tag{3}$$

##### 3.1. **Trivial steady state.**

We have the following result for the trivial steady state, where all microbial species/strains wash out of the chemostat.

**Theorem 3.1.** *In the absence of S. enterica, i.e when $S = 0$, the only steady state is a trivial steady state with $E_m = 0, E_r = 0$. At this steady state, the concentration of amino acids and acetate are zero $c = 0, a = 0, m = 0$ and the lactose is given by rates influx and dilution rates of the chemostat $\ell = \frac{L}{\gamma}$.*

*Proof.* Assume that $S = 0$, so there is no *S. enterica* present. Then from the last two equations in (3) we get $m = 0$ and $a = 0$. This forces in first two equations to have $E_m = E_r = 0$, which then forces $c = 0$ and, finally, $l\ell = \frac{L}{\gamma}$. This is the trivial steady state. □

**Theorem 3.2.** *The trivial steady state is asymptotically stable i.e. all solutions starting nearby will converge to it.*

The proof is delegated to section 4.

3.2. **Coexistence steady states.** Now we will investigate the existence of steady states where some of the microbial community members coexist. We start the analysis by looking for a steady state where all three members are present i.e. $E_m > 0, S > 0, E_r > 0$. With this assumption, it follows from the first three equations in (3) that

$$\mu_{Em} f(\ell) f(m) = \gamma \tag{4}$$

$$\mu_{Er} f(\ell) f(a) = \gamma \tag{5}$$

$$\mu_{Smr} f(c) = \gamma. \tag{6}$$

Using these relations in the last four equations in (3) we get

$$\begin{aligned} \ell &= -(E_m + E_r) + \frac{L}{\gamma} \\ c &= p_{Em,ac} E_m + p_{Er,ac} E_r - S \\ m &= -E_m + pS \\ a &= -E_r + pS. \end{aligned} \tag{7}$$

We first solve the equation (6). Since the function $f$ is increasing, there is a unique steady state value of acetate, which we denote $c^*$, which satisfies

$$\mu_{Smr} f(c^*) = \gamma.$$

From now on, we will denote steady state concentrations by asterisk $^*$. From the second equation in (7), we obtain a restriction on the steady state balance between consumption of acetate and the production of acetate $c^* = p_{Em,ac} E_m^* + p_{Er,ac} E_r^* - S^*$. We express steady state concentration $S^*$ as a function of steady state concentrations of both *E.* coli

$$S^* = p_{Em,ac} E_m^* + p_{Er,ac} E_r^* - c^*. \tag{8}$$

This population of *S. enterica* produces a steady state value of amino acids

$$m^* = -E_m^* + p(p_{Em,ac} E_m^* + p_{Er,ac} E_r^* - c^*) \tag{9}$$

$$a^* = -E_r^* + p(p_{Em,ac} E_m^* + p_{Er,ac} E_r^* - c^*) \tag{10}$$

The steady state value of lactose is also expressed as a function of $E_m^*, E_r^*$

$$\ell^* = \frac{L}{\gamma} - (E_m^* + E_r^*). \tag{11}$$

Any steady state of the system (2) has to satisfy equations (8, 9, 10, 11) with population concentrations $E_m^* \geq 0, E_r^* \geq 0$. The additional constraints come from the equations (4, 5, 6).

We now discuss different types of coexistence steady states.

3.2.1. *2-strain coexistence steady states.* We first discuss conditions for existence of a steady state $Q_r$ with

$$E_r > 0, S > 0, E_m = 0.$$

Only one strain of *E. coli* is present in a mutualistic coexistence with *S. enterica.*

**Theorem 3.3.** *The steady state $Q_r$, if it exists, is always unstable.*

*Proof.* Since $E_r > 0$, the equation (5) is valid at the steady state $Q_r$, which states that the growth rate of $E_r$ must be equal to the dilution rate of the chemostat. Using equations (11) and (9) with $E_m = 0$ we get

$$\mu_{Er} f(\frac{L}{\gamma} - E_r^*) f(-E_r^* + p(p_{Er,ac} E_r^* - c^*)) = \gamma.$$

This is a single equation in $E_r^*$ which may, or may not, have a positive solution. Note that the argument of the first function is the steady state amount of lactose that decreases with $E_r^*$ and the argument of the second function is the steady state concentration of methionine. If these values are sufficiently high to produce a growth rate for $E_r$ that matches the dilution rate, the two strain coexistence steady state is possible.

If steady state $Q_r$ exist, then at $Q_r$ the following holds

$$\begin{aligned} \mu_{Em} f(\ell^*) f(m^*) &= \mu_{Em} f(\frac{L}{\gamma} - E_r^*) f(p(p_{Er,ac} E_r^* - c^*)) \\ &> \mu_{Er} f(\frac{L}{\gamma} - E_r^*) f(p(p_{Er,ac} E_r^* - c^*)) \\ &> \mu_{Er} f(\frac{L}{\gamma} - E_r^*) f(-E_r^* + p(p_{Er,ac} E_r^* - c^*)) \\ &= \gamma. \end{aligned} \tag{12}$$

We show in Lemma 4.1 that one of the eigenvalues of the Jacobian $J(Q_r)$ is

$$\lambda = \mu_{Em} f(\ell^*) f(m^*) - \gamma.$$

By (12) it follows that $\lambda > 0$ and therefore $Q_r$, if it exists, is always unstable.

*Remark* 3.4. The positive eigenvalue $\lambda$ implies instability of $Q_r$ and is equivalent to a statement that the steady state $Q_r$ can be invaded by an infinitesimal population of $E_m$.

*Remark* 3.5. Note that the argument in (12) that shows $\lambda > 0$ depends on two inequalities. The first inequality is between the growth rates $\mu_m > \mu_r$; it is not surprising that the strain with the faster specific growth rate may invade a consortium of the slower growing *E. coli.* Importantly, the second inequality is based on the fact that the concentration of available methionine is greater than the concentration of available arginine, because the arginine is being consumed by $E_r$, but methionine, while being produced by *S. enterica*, is not being utilized at $Q_r$ where $E_m^* = 0$. Therefore, production of methionine by *S. enterica* is creating a niche that can be exploited by invading $E_m$. Crucially, as we show below, this type of niche created by faster growing $E_m$ can be exploited by invading *E. coli* $E_r$ with a slower specific growth rate.

We now discuss conditions for existence of a steady state $Q_m$ with

$$E_m > 0, S > 0, E_r = 0.$$

The argument is completely analogous with one important distinction. Note that only the first inequality in (12) depends on $\mu_m > \mu_r$, but the second does not. So it is possible that if the growth rates of the two *E. coli* strains are similar, as is the case for *E. coli* considered in this paper, then the slower growing strain $E_r$ will invade steady state $Q_m$.

This leads to the following result:

**Theorem 3.6.** *Assume that at the steady state $Q_m$, if it exists, we have*

$$\mu_{Er} f(a^*) > \mu_{Em} f(m^*). \tag{13}$$

*Then $Q_m$ is unstable and infinitesimal population of $E_r$ can invade $Q_m$.*

*Proof.* The argument is analogous to the proof of Theorem 3.3. The key argument, analogous to (12) is as follows

$$\mu_{Er} f(\ell^*) f(a^*) > \mu_{Em} f(\ell^*) f(m^*) = \gamma.$$

The inequality is the assumption (13) and the equality follows from the the equation (4) that holds at $Q_m$.

The instability now follows from Lemma 4.1 that shows that one of the eigenvalues of Jacobian $J(Q_m)$ is

$$\mu = \mu_{Er} f(\ell^*) f(a^*) - \gamma.$$

*Remark* 3.7. The assumption (13) states that if the excess of the unused available resource $a^*$ over utilized resource $m^*$ can overcome the slower specific growth rate $\mu_{Er} < \mu_{Em}$, the strain with the slower growth rate can invade a consortium of *E. coli* with faster growth rate.

##### 3.3. **Coexistence of 3 community members.**

Consider three species/strain steady state $C$ (for 'coexistence')

$$E_m \neq 0, S \neq 0, E_r \neq 0.$$

The equations (4,5) together with (9,10,11) give

$$\mu_{Em} f(\frac{L}{\gamma} - E_m^* - E_r^*) f(-E_m^* + p(p_{Em,ac}E_m^* + p_{Er,ac}E_r^* - c^*) = \gamma$$

$$\mu_{Er} f(\frac{L}{\gamma} - E_m^* - E_r^*) f(-E_r^* + p(p_{Em,ac}E_m^* + p_{Er,ac}E_r^* - c^*)) = \gamma.$$

This system of nonlinear equations in $E_m^*, E_r^*$ may, or may not, have a positive solution. This depends delicately on the values of parameters and we were unable to find general relationships between the parameters that would guarantee existence of such a solution. When $C$ does not exist and $Q_r, Q_m$ are unstable, the population is not sustainable and dies out by converging to the trivial steady state.

However, the formulas above imply that the following parameter changes increase the steady state growth rates of $E_m$ and $E_r$ on the right hand side of the equations and therefore make it easier for these strains of *E. coli* to avoid washing out of the chemostat. As such, these changes promote existence of coexistence steady state $C$:

- simultaneous increases in specific growth rates $\mu_{Em}, \mu_{Er}$;
- increase in lactose supply $L$, or a decrease in dilution rate $\gamma$;
- increase in acetate production yields $p_{Em,ac}, p_{Er,ac}$;
- simultaneous increase in amino acid production yields $p = p_{Smr,met} = p_{Smr,arg}$;

- an increase in specific growth rate $\mu_{Smr}$ will make steady state value of $c^*$ smaller. This, in turn, will increase the argument of the second functions in both equations, making the right hand side larger.

We provide numerical simulation for parameters examined in the main body in Section 5.

#### 4. Stability of the equilibria.

To determine stability of the equilibria, we compute a Jacobian $J$ of the system (2):

$$\begin{bmatrix} \mu_m f(\ell)f(m)-\gamma & 0 & 0 & E_m\mu_m f(m)f'(\ell) & 0 & E_m\mu_m f'(m)f(\ell) & 0 \\ 0 & \mu_r f(\ell)f(a)-\gamma & 0 & E_r\mu_r f(a)f'(\ell) & 0 & 0 & E_r\mu_r f'(a)f(\ell) \\ 0 & 0 & \mu_S f(c)-\gamma & 0 & S\mu_S f'(c) & 0 & 0 \\ -\mu_m f(\ell)f(m) & -\mu_r f(\ell)f(a) & 0 & A & 0 & -E_m\mu_m f'(m)f(\ell) & -E_r\mu_r f'(a)f(\ell) \\ p_m\mu_m f(\ell)g(m) & p_r\mu_r f(\ell)h(a) & \mu_S f(c) & B & -\mu_S S f'(c)-\gamma & p_m E_m\mu_m f'(m)f(\ell) & p_r E_r\mu_r f'(a)f(\ell) \\ -\mu_m f(\ell)f(m) & 0 & p\mu_S f(c) & -E_m\mu_m f(m)f'(\ell) & p\mu_S S f'(c) & -E_m\mu_m f'(m)f(\ell)-\gamma & 0 \\ 0 & -\mu_r f(\ell)f(a) & p\mu_S f(c) & -E_r\mu_r f(a)f'(\ell) & p\mu_S S f'(c) & 0 & -E_r\mu_r f'(a)f(\ell)-\gamma \end{bmatrix}.$$

where we use the following notation in order to fit the matrix to the page

$$\begin{aligned} A &:= -f'(\ell)(E_m\mu_m f(m)+E_r\mu_r f(a))-\gamma \\ B &:= f'(\ell)(p_m E_m\mu_m f(m)+p_r E_r\mu_r f(a)) \\ p_m &= p_{Em,ac}, p_r = p_{Er,ac}, \\ \mu_m &= \mu_{Em}, \mu_r = \mu_{Er}, \mu_S = \mu_{Smr}. \end{aligned}$$

**Proof of Theorem 3.2**

We evaluate the Jacobian at the trivial steady state to get

$$\begin{bmatrix} -\gamma & 0 & 0 & 0 & 0 & 0 & 0 \\ 0 & -\gamma & 0 & & 0 & 0 & 0 \\ 0 & 0 & -\gamma & 0 & 0 & 0 & 0 \\ 0 & 0 & 0 & -\gamma & 0 & 0 & 0 \\ 0 & 0 & 0 & 0 & -\gamma & 0 & 0 \\ 0 & 0 & 0 & 0 & 0 & -\gamma & 0 \\ 0 & 0 & 0 & 0 & 0 & 0 & -\gamma \end{bmatrix}.$$

This matrix has 7 eigenvalues with negative real part and hence is asymptotically stable.

**Lemma 4.1.** *The Jacobian evaluated at the two species steady state $Q_r$ has an eigenvalue*

$$\lambda = \mu_{Em} f(\ell^*) f(m^*) - \gamma.$$

*The Jacobian evaluated at the two species steady state $Q_m$ has an eigenvalue*

$$\mu = \mu_{Er} f(\ell^*) f(a^*) - \gamma$$

*Proof.* We evaluate the Jacobian at an steady state $Q_r$ with $E_m = 0, S > 0, E_r > 0$ which implies

$$\mu_{Smr} f(c) = \gamma \quad \text{and} \quad \mu_{Er} f(\ell) f(a) = \gamma.$$

The Jacobian becomes

$$\begin{bmatrix} \mu_{Em} f(\ell) f(m) - \gamma & 0 & 0 & 0 & 0 & 0 & 0 \\ 0 & 0 & 0 & E_r \mu_{Er} f(a) f'(\ell) & 0 & 0 & E_r \mu_{Er} f'(a) f(\ell) \\ 0 & 0 & 0 & 0 & S \mu_{Smr} f'(c) & 0 & 0 \\ -\mu_{Em} f(\ell) f(m) & -\gamma & 0 & -f'(\ell) E_r \mu_{Er} f(a) - \gamma & 0 & 0 & -E_r \mu_{Er} f'(a) f(\ell) \\ p_m \mu_{Em} f(\ell) f(m) & p_r \gamma & -\gamma & f'(\ell) p_r E_r \mu_{Er} f(a)) & -\mu_{Smr} S f'(c) - \gamma & 0 & p_r E_r \mu_{Er} f'(a) f(\ell) \\ -\mu_{Em} f(\ell) f(m) & 0 & p\gamma & 0 & p \mu_{Smr} S f'(c) & -\gamma & 0 \\ 0 & -\mu_{Er} f(\ell) f(a) & p\gamma & -E_r \mu_{Er} f(a) f'(\ell) & p \mu_{Smr} S f'(c) & 0 & -E_r \mu_{Er} f'(a) f(\ell) - \gamma \end{bmatrix}.$$

We note that by expanding the Jacobian along the first row, one of the eigenvalues is

$$\lambda = \mu_{Er} f(\ell^*) f(a^*) - \gamma.$$

An analogous argument shows that the Jacobian at $Q_m$, where

$$\mu_{Smr} f(c) = \gamma \quad \text{and} \quad \mu_{Em} f(\ell) f(m) = \gamma$$

has an eigenvalue

$$\mu = \mu_{Er} f(\ell^*) f(a^*) - \gamma.$$

#### 5. Numerical simulations

We illustrate the existence of three community member coexistence steady states, where both strains of *E. coli* coexist with *S. enterica*, by numerical simulation using the same parameters that were used in the batch model. Since we are using a chemostat model, there are two new parameters: $L$, which represents the concentration of lactose in the influx to the chemostat and the dilution rate $\gamma$. For the simulations, we will use parameters 1 and

$$L = 0.8, \qquad \gamma = 0.4.$$

The simulations show that there are two coexistence steady states, one of which is locally asymptotically stable, denoted $C$ for "coexistence", and the other which is a saddle point, denoted $S$.

$$S := (E_m, E_r, Smr, lcts, ac, met, arg) = (0.3198, 0.3189, 0.3256, 1.6345, 0.3132, 0.0058, 0.0067)$$
$$C := (E_m, E_r, Smr, lcts, ac, met, arg) = (1.8571, 0.1362, 1.9535, 0.0067, 0.0400, 0.0966, 1.8175)$$

The saddle point is unstable with one-dimensional unstable manifold. The stability properties have been confirmed by evaluating the Jacobian at $C$ and $S$ and computing eigenvalues.

We summarize our observations in Figure 1. In the left panel, we show a conceptual picture in the phase space $\mathbf{R}^7$, which shows bistability between the trivial washout steady state and the coexistence steady state $C$. Basins of attractions of these points are separated by a six dimensional stable manifold of the saddle point $S$. Therefore, which steady state the population will reach will depend on the initial population and the initial distribution of resources. In the right panel, we project three trajectories starting at three different initial conditions to a two dimensional space of $E_m, E_r$ concentrations. The red dashed line with initial condition

$$(E_m, E_r, Smr, lcts, ac, met, arg) = (1, 0.1, 1, 10, 1, 1, 1)$$

and the blue dashed-dot line initial condition

$$(E_m, E_r, Smr, lcts, ac, met, arg) = (0.1, 1, 1, 10, 1, 1, 1)$$

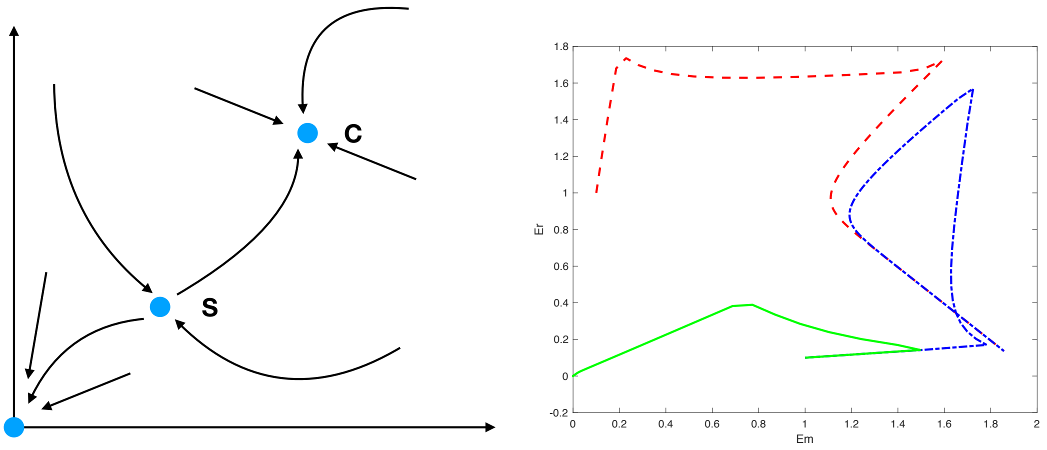


FIGURE 1. (Left) Conceptual picture of the 7-dimensional phase space. The trivial steady state is stable. If the coexistence steady states exist, then $S$ is a saddle and $C$ is stable. (Right) Projections of three trajectories in $\mathbf{R}^7$ to into $\mathbf{R}^2$ of concentrations of two *E. coli* strains. The red dashed line with Initial condition $(E_m, E_r, Smr, lcts, ac, met, arg) = (1, 0.1, 1, 10, 1, 1, 1)$ and the blue dashed-dot line initial condition $(0.1, 1, 1, 10, 1, 1, 1)$ both converge to coexistence steady state $C$ in the lower right of the Figure. The green line curve starts at the same values of $E_m, E_r$ as the blue curve, but with a small initial concentration of *S. enterica* $(1, 0.1, 0.01, 10, 1, 1, 1)$; this population converges to the trivial steady state and does not survive.

both converge to coexistence steady state $C$ in the lower right corner of Figure 1. The green line starts at the same values of $E_m, E_r$ as the blue curve, but with a small initial concentration of *S. enterica*

$$(E_m, E_r, Smr, lcts, ac, met, arg) = (1, 0.1, 0.01, 10, 1, 1, 1);$$

this population converges to the trivial steady state and does not survive.

We illustrate the dynamics changes in the consortium composition along the way to coexistence steady state $C$ in Figure 2. We start with initial values

$$(E_m, E_r, Smr, lcts, ac, met, arg) = (1, 1, 1, 1000, 1, 1, 1)$$

and graph the behavior of all microbial strains and resources, except lactose. The lactose initial concentration is high to match the batch experiments and its inclusion skews the axis in a way that obscures the dynamic periodic changes in the population dynamics. We observe fast exponential growth in $E_r$ and $E_m$ strains until both methionine and arginine are exhausted; the acetate that the *E. coli* produce allows the slower growing *S. enterica* to continue to grow and thus replenish the amino acid pools in the chemostat. This induces a rebound in the *E. coli* strains around time 12. The exhaustion of the acetate pool leads to a decline of the *S. enterica* population toward the steady state, starting around the time 15. This decline induces a second decline in *E. coli* populations, which then slowly converge to their steady state values. Note that the steady state value of $E_m$

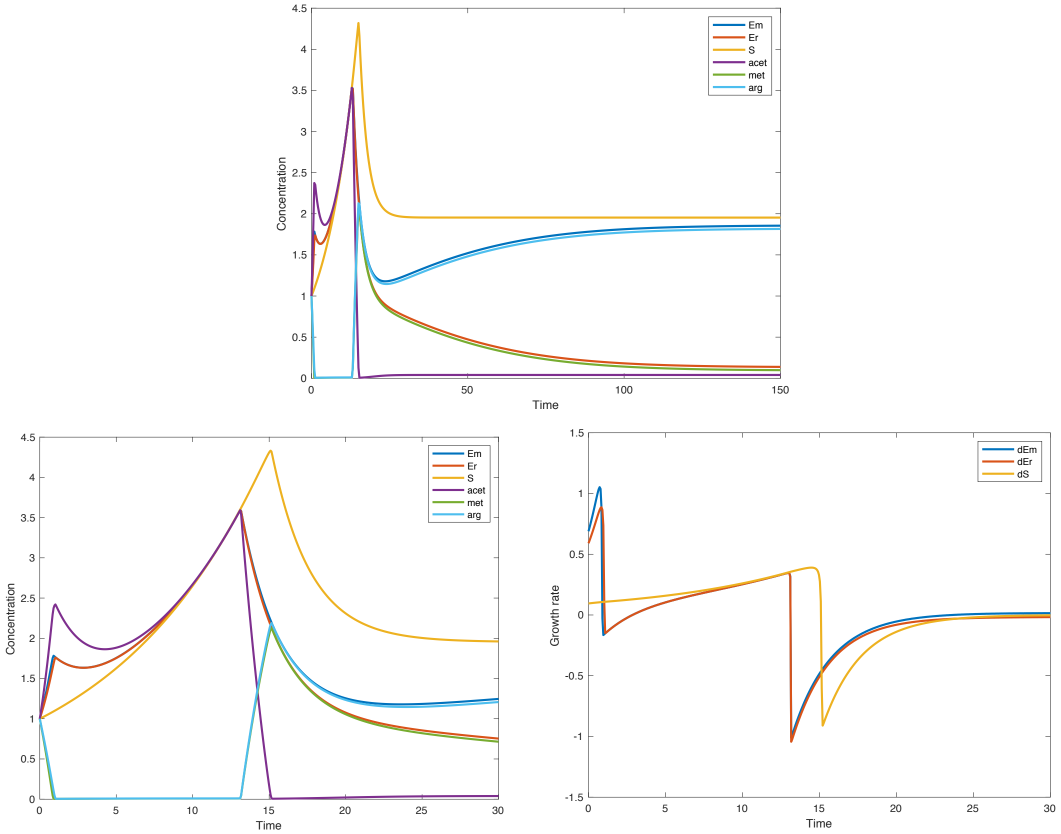


Figure 2. (Top) Concentrations converge to the stable steady state $C$ for the parameters listed above; (Lower Left) Detail of the concentrations for first 30 units of time. (Lower Right) Growth rates of the *E. coli* strains and *S. enterica* show rapid changes along the way to the steady state. Initial data for all three figures $(E_m, E_r, Smr, lcts, ac, met, arg) = (1, 1, 1, 1000, 1, 1, 1)$.

is higher than that of $E_r$, reflecting the difference in higher specific growth rates between these strains.
